# Structural Basis of TACO1-Mediated Efficient Mitochondrial Translation

**DOI:** 10.64898/2025.12.18.695269

**Authors:** Shuhui Wang, Michele Brischigliaro, Yuekang Zhang, Chunxiang Wu, Wei Zheng, Antoni Barrientos, Yong Xiong

**Author notes:** These authors contributed equally to this work (Shuhui Wang, Michele Brischigliaro). Corresponding author. (A.B.); (Y.X.).

## Abstract

Translation elongation is a universally conserved step in protein synthesis, relying on elongation factors that engage the ribosomal L7/L12 stalk to mediate aminoacyl-tRNA delivery, accommodation, and ribosomal translocation. Using *in organello* cryo-electron microscopy, we reveal how the mitochondrial translation accelerator TACO1 promotes efficient elongation on human mitoribosomes. TACO1 binds the mitoribosomal region typically bound by elongation factor Tu (mtEF-Tu), bridging the large and small subunits via contacts with 16S rRNA, bL12m, A-site tRNA, and uS12m. While active throughout elongation, TACO1 is especially critical when translating polyproline motifs. Its absence prolongs mtEF-Tu residence in A/T states, causes persistent mitoribosomal stalling and premature subunit dissociation. Structural analyses indicate that TACO1 competes with mtEF-Tu for mitoribosome binding, stabilizes A-site tRNA, and enhances peptidyl transfer through a mechanism distinct from EF-P and eIF5A. These findings suggest that bacterial TACO1 orthologs may serve analogous roles, highlighting an evolutionarily conserved strategy for maintaining elongation efficiency during challenging translation events.

## Introduction

Translation elongation is a central and highly regulated phase of protein synthesis, during which ribosomes decode mRNA into polypeptide chains with speed and precision. This stage is crucial for shaping the proteome and enabling rapid cellular responses to physiological demands^1^. While elongation dynamics have been extensively characterized in bacterial and cytosolic translation systems^2,3^, their regulation in mitochondria is only starting to emerge^4^.

Mitoribosome, a dedicated translational machinery in mitochondria, is responsible for synthesizing mitochondrial DNA (mtDNA)-encoded core components of the oxidative phosphorylation (OXPHOS) system that powers cellular aerobic energy production^5^. Although descended from bacterial ribosomes, mitoribosomes have undergone extensive remodeling in composition and structure^6,7^. In mammals, the 55S mitoribosome comprises a 28S small subunit (mtSSU) and a 39S large subunit (mtLSU), composed of 82 proteins and three ribosomal RNAs (rRNAs). In addition to the 12S rRNA in the mtSSU and the 16S rRNA in the mtLSU, a structural tRNA^Val/Phe^ replaces the canonical 5S rRNA^8–10^, and overall rRNA content is markedly reduced (∼25–30%) compared to bacterial and cytosolic ribosomes (∼65%), reflecting major architectural remodeling during evolution^11^.

Accurate and efficient mitochondrial translation is critical for OXPHOS biogenesis, and its disruption contributes to diverse neurometabolic and systemic diseases, including Leigh syndrome, sensorineural hearing loss, and hypertrophic cardiomyopathy^12–14^. These pathologies often stem from mutations in mitochondrial tRNAs, rRNAs, mitoribosomal proteins, or translation factors^15,16^. Among the latter, mutations in human TACO1 (Translation Accelerator of Cytochrome *c* Oxidase subunit 1-CO1) cause complex IV deficiency and Leigh syndrome^17–22^. We recently showed that TACO1 alleviates mitoribosomal stalling at polyproline stretches, particularly at a unique triple-proline motif in CO1^23^, highlighting how nascent chain sequences influence elongation.

In prokaryotes, stalling at polyproline tracts is resolved by elongation factor P (EF-P)^24–26^, and members of the ATP-binding cassette family-F (ABCF)^27–29^, whereas in eukaryotes, initiation factor 5A (eIF5A) performs this function^30^. Despite structural divergence, these factors converge on a common mechanism: binding the ribosomal E-site to stabilize peptidyl-tRNA in the P-site and promote peptide bond formation^31–37^. However, no direct homologs of these factors have been identified in mitochondria, and the mechanism by which TACO1 mitigates stalling remains unclear. Biochemical studies suggest TACO1 forms a labile ∼74 kDa complex with mitochondrial translation factors^22^ and a small fraction associate with the mtSSU^38^, mtLSU and the monosome, potentially near the tRNA-acceptor site (A-site) ^23^, but its precise mode of action remains elusive.

Recent advances in cryo-electron microscopy (cryo-EM) have revealed key features of isolated mitoribosomes^39–47^, but these *in vitro* structures may miss key conformational states and interactions with mitochondrial membranes or transient regulatory factors. *In situ* cryo-electron tomography (cryo-ET) has shown that mitoribosomes are tethered to the mitochondrial inner membrane via by the mtLSU protein mL45^48,49^, aligning the polypeptide exit tunnel with the OXA1L insertase for co-translational membrane insertion^50–53^. Nonetheless, achieving near-atomic resolution of mitoribosomes in mitochondria has been limited by their low abundance and poor signal-to-noise ratios.

In this study, we present *in oganello* cryo-EM structures of human mitoribosomes tethered to the inner mitochondrial membrane, reaching 2.5 Å resolution. We identify TACO1 as a mitoribosome-associated factor that competes with elongation factor Tu (mtEF-Tu) for mitoribosome binding, stabilizing A-site tRNA, and contributing to peptidyl transfer. Phylogenetic analyses show conservation of TACO1 orthologs in prokaryotes^54,55^ and mitochondria, highlighting an ancient mechanism for managing challenging elongation events, regulating translational elongation, and enhancing translational efficiency. Our findings uncover a fundamental strategy for safeguarding mitochondrial protein synthesis and enhancing translational efficiency.

## Results

### Structure of the human mitoribosome within isolated mitochondria

To capture the mitoribosome structure in an environment as close to native as possible, intact mitochondria were isolated from human cells using gradient centrifugation and prepared for single particle cryo-EM data collection (Fig. 1a). The thickness of the mitochondrial samples, combined with the lower nucleotide content of mitoribosomes compared to bacterial and cytosolic ribosomes^56–58^, result in significantly reduced signal-to-noise ratios in cryo-EM micrographs, complicating particle identification (Extended Data Fig. 1a). To overcome this, 2D template matching (2DTM)^56^ and GisSPA^59^ were employed for particle picking and initial alignment, using a template derived from a previously published human mitoribosome structure^39^ (Extended Data Fig. 1b). To minimize reference bias, the template was modified prior to particle picking by removing a helix from the 16S rRNA and applying lowpass filtering to 8 Å resolution (Extended Data Fig. 1b). Following 3D classification, particles restoring the deleted helix were retained, yielding a consensus reconstruction at 2.5 Å resolution—a striking improvement over the reference map (Extended Data Fig. 1d and Extended Data Tables 1). Focused classification on the tRNA sites further separated the biologically relevant functional states of the mitoribosome, each resolved at ∼2.9-3.1 Å resolution (Extended Data Fig. 1e and Extended Data Tables 1-2).

**Fig. 1.**
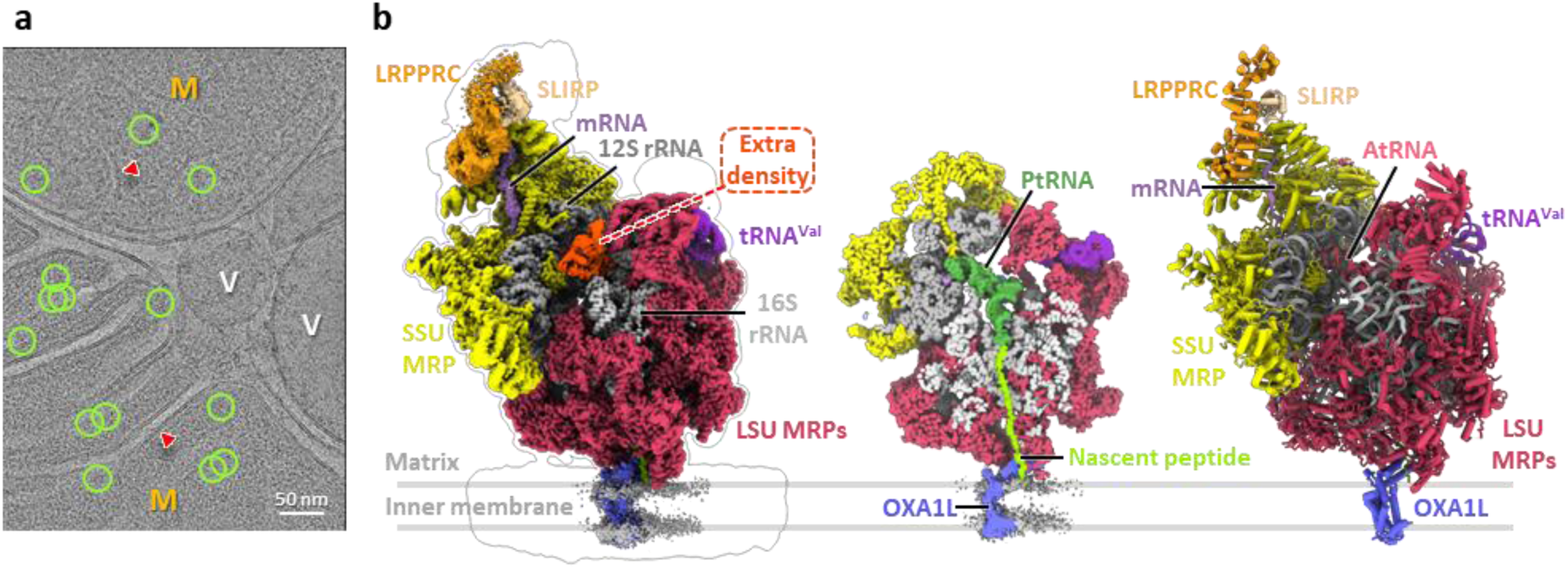
| Cryo-EM structure of human mitoribosome within mitochondria. (**a**) Representative cryo-EM micrograph of human mitochondria. M, mitochondria; V, vesicle. Red arrowheads mark cristae, and green circles mark the final particles used for reconstruction of the 55S mitoribosome mapped-back onto the micrograph. (**b**) Cryo-EM density map of the mitoribosome tethered to the inner membrane (left), a cross-sectional view through the nascent peptide tunnel (middle), and cartoon representation of the corresponding atomic model (right).

This high-resolution map allowed the identification of clear side-chain features and base-pairing interactions, enabling precise atomic modeling (Fig. 1b, Extended Data Fig. 2a and 2b). rRNA modifications and cofactors, including adenosine triphosphate (ATP), nicotinamide adenine dinucleotide (NAD), and spermine (SPM), were unambiguously identified, with their structural models fitting well into the corresponding densities (Extended Data Fig. 2b), consistent with the highest-resolution in vitro structure of the human mitoribosome^39^. Notably, in contrast with the reference map used for particle picking, we observed an additional density by the end of the polypeptide exit tunnel (Fig. 1b). This density was identified as the OXA1L-mitoribosome interaction region that facilitate co-translational membrane insertion^50–53^, and the docking of an AlphaFold-predicted model of OXA1L^60^ into this density showed good agreement with the density (Extended Data Fig. 3). Interestingly, unlike bacterial and cytosolic ribosomes, all reconstructed 55S mitoribosome particles in our dataset appeared to be membrane-associated, with no free 55S mitoribosome observed. This finding is consistent with cryo-ET studies reporting that ∼80% of mitoribosomes in yeast and human mitochondria reside within 30 nm of the inner mitochondrial membrane^48,49^ and that *Chlamydomonas reinhardtii* mitoribosomes are exclusively tethered to the inner membrane^61^. Together with systematic mutant analysis in yeast^62^, these results suggest that the mtLSU assembles onto the membrane prior to forming the 55S initiation complex with the mtSSU. Compared to the cryo-ET map of *Chlamydomonas* mitoribosomes^61^, we did not observe a second membrane contact site in human mitoribosome. Additionally, aligning with previous cryo-ET findings^49^, we observed another additional density near the mRNA entry site on the mtSSU (Fig. 1b), absent in the reference map. This density was identified as the mRNA delivery complex LRPPRC-SLIRP^63^.

**Fig. 2.**
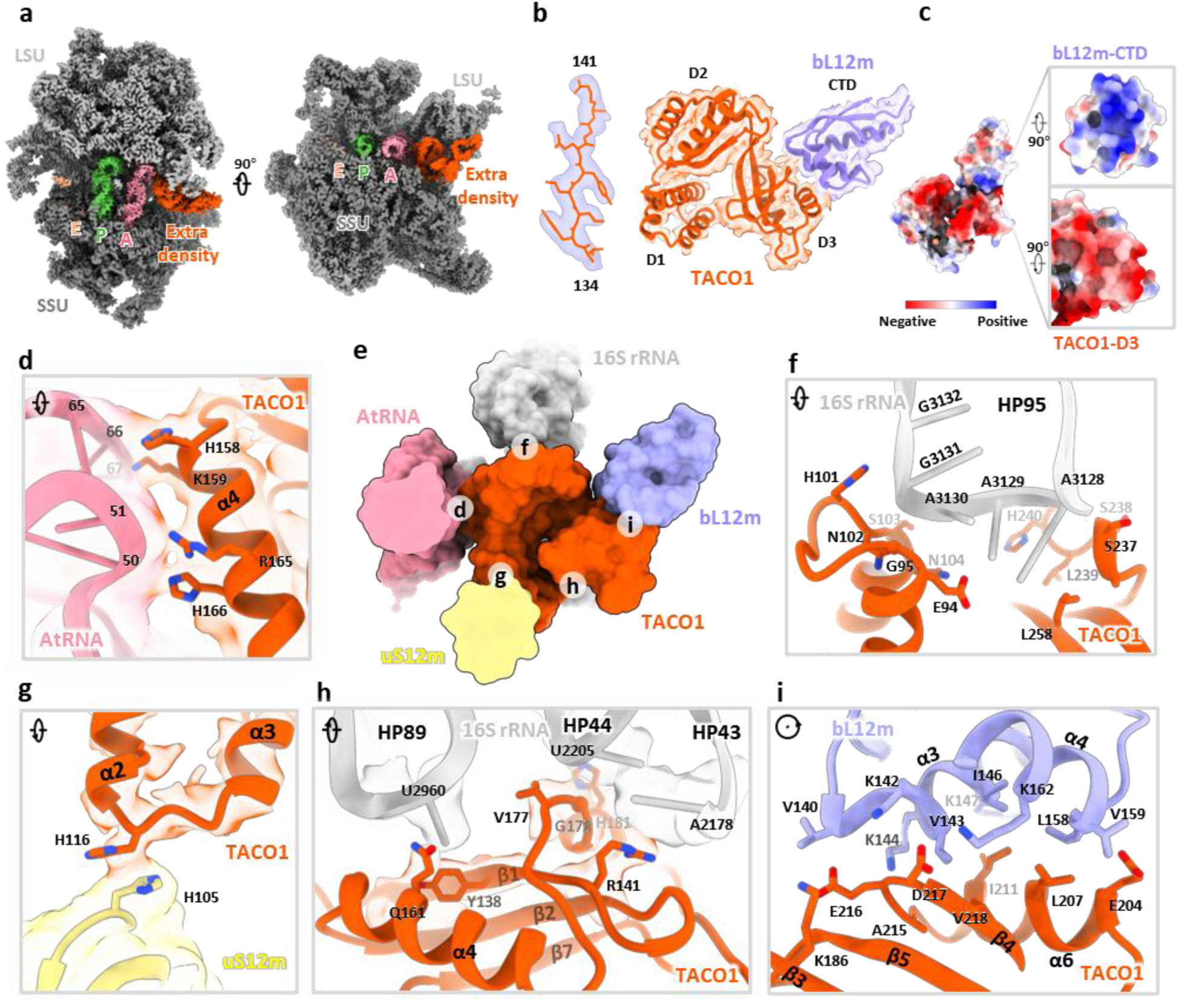
| Structural identification of the translation factor TACO1 and its interactions with the mitoribosome. (**a**) Cross-sectional cryo-EM maps of the mitochondrial ribosome, sectioned through the tRNAs-binding region, show an unassigned density adjacent to the A-site tRNA. (**b**) Focused views of the extra density fitted with a portion of TACO1 (left) and with combined TACO1 and bL12m models (right). (**c**) Electrostatic surface potential at the TACO1-bL12m interface, highlighting complementary charge distributions. (**d-i**) Detailed interaction network between TACO1 and mitoribosomal components (**e**), including direct contacts with the A-site tRNA (**d**), 16S rRNA (**f**, **h**), uS12m (**g**), and bL12m (**i**).

### Identification of TACO1 binding on mitoribosome

Beyond recapitulating previously reported features, we identified an unassigned density adjacent to the A-site tRNA, a feature not captured in any earlier *in vitro* studies (Fig. 2a). To resolve this unknown density, we performed local refinement, improving the resolution to 2.4 - 4 Å across the unassigned region (Extended Data Fig. 4b). Partial main-chain tracing, followed by DALI-based structural comparison^64^ against the AlphaFold database, identified the translation factor TACO1 as the top candidate (Extended Data Fig. 4a). Fitting a complete atomic model of TACO1 into the density map revealed an excellent match, supporting its unambiguous assignment (Fig. 2b and Extended Data Fig. 4d). Prior studies using sucrose gradients and immunoprecipitation had shown that a small fraction of mouse or human TACO1 associates with the mitoribosome^23,38^, while the structural footprint of TACO1 on the mitoribosome had remained unknown. Furthermore, we resolved the C-terminal domain of the mitoribosomal protein bL12m (bL12m-CTD) in the remaining density, positioned in direct contact with TACO1 (Extended Data Fig. 4a). The bL12m-CTD lies within the expected reach of its flexible linker connecting to the N-terminal domain (NTD), which is anchored on the L7/L12 stalk^39^, consistent with prior BioID data identifying bL12m as the primary proximal interactor of TACO1^23^. At the binding interface, electrostatic complementarity is evident: the bL12m surface is predominantly positively charged, featuring residues K142, K144, K147, and K162, while TACO1 presents a negatively charged patch composed of E216, D217, and E204 (Fig. 2, c and i), contributing to the electrostatic interaction between the two proteins.

Notably, TACO1 engages not just with bL12m, but also establishes extensive contacts with key mitoribosomal components: the A-site tRNA, 16S rRNA on the mtLSU, and uS12m on the mtSSU. Positively charged residues, R165, H166, H158 and K159, in TACO1 helix 4 (α4) extend toward and interact with the stem region of the A-site tRNA (Fig. 2d). Additionally, H116, located in the loop between α-helices 2 (α2) and 3 (α3) of TACO1 forms a stacking contact with H105 in uS12m (Fig. 2g). Moreover, TACO1 engages extensively with the loop regions of 16S rRNA hairpins 43 (HP43), 44 (HP44), 89 (HP89), and 95 (HP95) (Fig. 2, f and h). These extensive, multivalent contacts suggest that TACO1 is optimally positioned to stabilize the elongation complex during critical phases of mitochondrial translation.

### Competitive binding of TACO1 and mtEF-Tu on the mitoribosome

TACO1 was initially identified through clinical studies linking its loss-of-function mutations to impaired *CO1* mRNA translation, respiratory complex IV deficiency, and a late-onset form of Leigh syndrome^22^. Five pathogenic mutations have been reported leading to translational frameshift, and premature termination at protein residues C85, R141, H158, E226 and E259, resulting in complex IV deficiency (Extended Data Table 5)^17–21^. Recently, we showed that TACO1 resolves polyproline-induced pauses not only during synthesis of CO1, but for all 13 mtDNA-encoded polypeptides^23^. However, as the only triple-proline motif in the mtDNA proteome occurs in CO1, its synthesis is most prominently affected, leading to a complex IV assembly defect^23^. These findings underscore the essential role of TACO1 in maintaining mitochondrial translation efficiency.

To gain mechanistic insight into the functional role of TACO1, we compared cryo-EM structures of mitoribosomes bound to TACO1 and to elongation factor Tu (mtEF-Tu) (Fig. 3a)^65^. Structural superposition revealed that although the TACO1-binding site partially overlaps with that of mtEF-Tu, TACO1 establishes a more extensive interface with the mitoribosome (Fig. 3b). While mtEF-Tu can only deliver aminoacyl-tRNAs to the A/T site, as it is sterically blocked by hairpin HP95 of the 16S rRNA, TACO1 binds HP95 and extends additional interactions to HP43, HP44 and HP89 to directly stabilize the A-site tRNA (Fig. 3, a and c). Further supporting a competitive binding model, our cryo-EM reconstructions of free 39S subunits revealed two distinct classes: one associated with TACO1, and another with an inactive conformation of mtEF-Tu (Extended Data Fig. 5), reminiscent of a previously reported late-maturation-state of the 39S mtLSU^47^. Structural superposition of these two states indeed revealed steric clashes between TACO1 and the inactive mtEF-Tu at their respective binding sites (Extended Data Fig. 6).

**Fig. 3.**
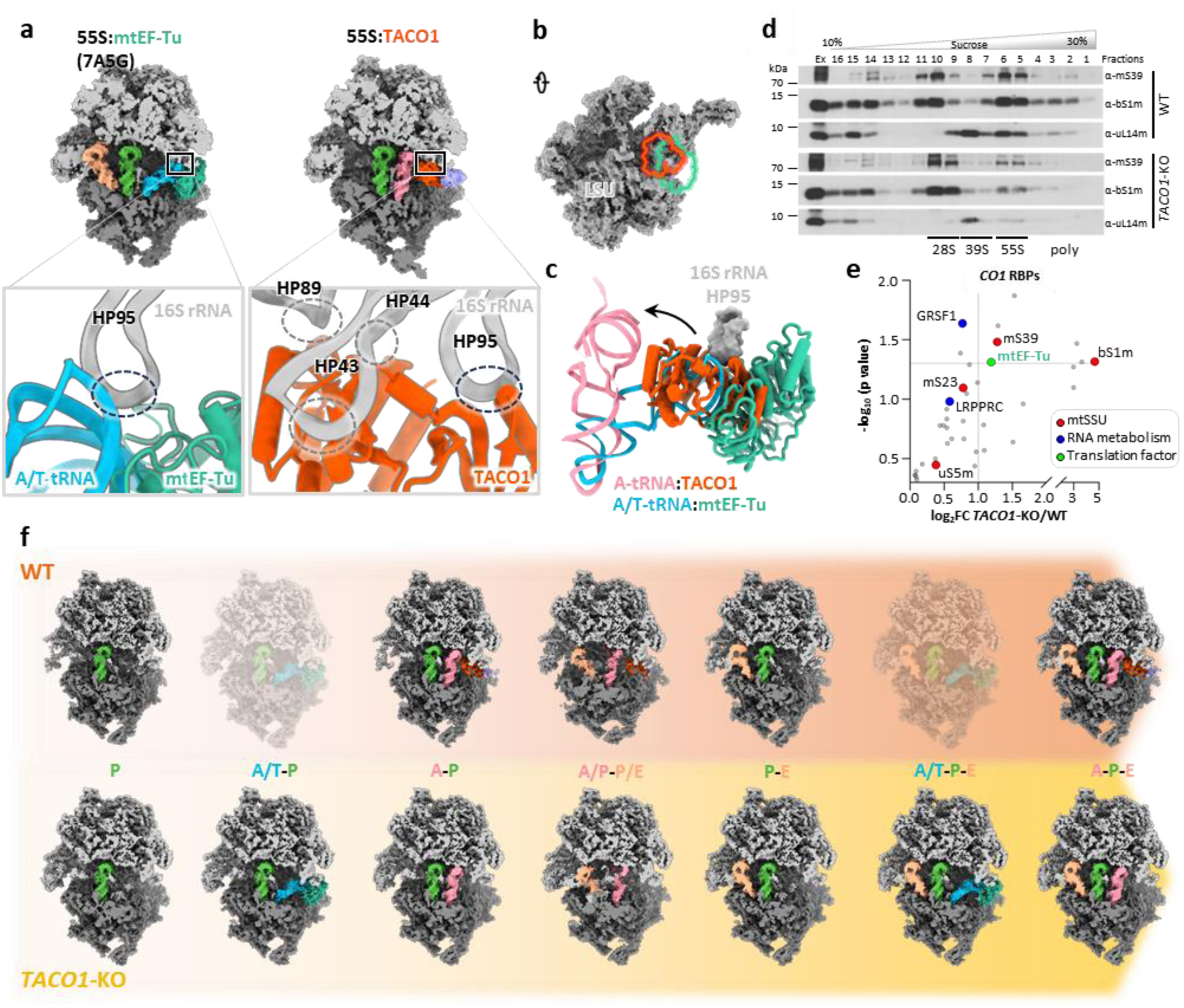
| Competitive binding of TACO1 and mtEF-Tu to mitoribosome in WT and *TACO1*-KO cells. (**a**) Binding sites of mtEF-Tu (left, PDBID: 7A5G) and TACO1 (right) on the 55S mitoribosome, with bottom panels highlighting their respective interaction surfaces (teal-green for mtEF-Tu and orange for TACO1) on the mtLSU. (**b**) Illustration of TACO1 (orange) and mtEF-Tu (teal green) binding sites on the mtLSU. (**c**) Structural superposition showing significant overlap between the binding sites of mtEF-Tu with A/T-tRNA, positioned below the hairpin 95 (HP95) of 16S mt-rRNA (gray), and TACO1, which binds beneath HP95 and displaces the tRNA toward the A site. (**d**) Sucrose gradient sedimentation profiles of mtSSU (mS39 and bS1m) and mtLSU markers (uL14m) in mitochondria purified from WT and *TACO1*-KO cells. (**e**) Volcano plot of proteins with increased *CO1* mRNA binding in *TACO1*-KO versus WT cells. Canonical RNA-binding, ribosome proteins and translation factors are highlighted and color coded. n = 3 biological replicates; dashed lines denote significance thresholds (adjusted p < 0.05, fold change > 2). (**f**) Sequential states of the mitoribosome in wild-type (WT, top) and *TACO1*-KO (bottom) human cell lines, highlighting the impact of TACO1 loss.

Given that mtEF-Tu functions as a GTPase^66^, we next tested whether recombinant TACO1 could directly bind GTP. This hypothesis was supported by three independent lines of evidence—biochemical analyses of native complexes^22^, Bio-ID proximity-labeling studies^23^ and in silico protein-protein interaction predictions^67^—all identifying high-confidence interactions between TACO1 and the guanine-nucleotide exchange factor mtEF-Ts. Using BODIPY-FL-GTPγS binding assays, we observed a robust increase in fluorescence upon TACO1 addition, consistent with direct GTP binding. This effect was abolished by pre-incubation with the non-hydrolyzable GTP analog GMPPNP, but not with its adenine-nucleotide counterpart AMPPNP (Extended Data Fig. 7a), indicating selectivity for guanine-nucleotides. The dissociation constant (K_d_) for TACO1 binding to BODIPY-FL-GTPγS was determined to be 25 μM (Extended Data Fig. 7b), similar to that reported for mtEF-Tu.GTP in vitro^66^. Together, these results, along with the observed structural overlap between the binding sites of TACO1 and the GTPase mtEF-Tu on both the 55S and 39S mitoribosomes, are consistent with a potential GTP-binding role for TACO1.

### TACO1 knockout leads to the accumulation of mitoribosome A/T states with mtEF-Tu bound

Immunoblot analyses of purified mitochondria indicated that the steady-state levels of mitoribosome components and mtEF-Tu were comparable between wild-type (WT) and *TACO1* knockout (*TACO1*-KO) samples (Extended Data Fig. 7c). To further investigate the role of TACO1 in mitochondrial translation, we analyzed in the two cell lines mitoribosome sedimentation profiles in sucrose gradients. Gradient fractions were analyzed by immunoblotting with antibodies against markers of the mtSSU (mS39 and bS1m) and the mtLSU (uL14m) (Fig. 3d). In WT cells, the levels of free mtSSU were comparable to those of the 55S monosome. In contrast, in *TACO1*-KO cells, free mtSSU levels were approximately twice those of the 55S monosome (Extended Data Fig. 7f). Similarly, in WT cells, free mtLSU levels were close to those of the 55S monosome, while in *TACO1*-KO cells, free mtLSU levels were approximately three times higher (Extended Data Fig. 7f). This shift indicates that TACO1 plays a critical role in stabilizing the 55S mitoribosome and preventing subunit dissociation.

Given the essential role TACO1 plays in *CO1* mRNA translation^22,23^, we next performed LC-MS/MS following UV-induced crosslinking to identify RNA-binding proteins enriched on *CO1* mRNA (Extended Data Fig. 7d). qPCR analysis confirmed specific enrichment of *CO1* mRNA. and the proportion of captured *CO1* mRNA relative to input was comparable between WT and *TACO1*-KO cells (Extended Data Fig. 7e). Canonical mRNA binding proteins such as LRPPRC and GRSF1 were captured with similar enrichment in both WT and *TACO1*-KO cells (Fig. 3e). In addition, the main mtSSU components of the mRNA channel (uS5m and mS39) were also pulled-down, reflecting effective capture of mitoribosome bound, actively translated *CO1* mRNA species (Fig. 3e). Notably, mtEF-Tu levels in this assay were significantly elevated in *TACO1*-KO cells relative to wild-type controls (Fig. 3e), implying prolonged mtEF-Tu residence on mitoribosomes when TACO1 is absent. This observation aligns with reported evidence that bacterial EF-Tu and eukaryotic eEF2 can be found associated with stalled ribosomal complexes^68–70^.

*In organello* Cryo-EM analysis of mitochondria from *TACO1*-KO cells yielded a 2.8 Å consensus mitoribosome map and resolved multiple elongation intermediates (Extended Data Fig. 8). Notably, unlike in WT cells, where the predominant elongation state was A-P (Extended Data Fig. 1e and Extended Data Fig. 7g), *TACO1*-KO mitoribosomes showed a clear enrichment of mtEF-Tu–bound A/T-P and A/T-P-E states that were not observed in WT cells (Fig. 3f and Extended Data Fig. 8c). Conversely, the A/T-P-E state was the most abundant species in *TACO1*-KO cells underscoring a defect in the transition from A/T to A states (Extended Data Fig. 7g). Comparison of density at the peptidyl transfer center further revealed that in the WT A-P-E state, the nascent peptide connects to the A-site tRNA, whereas in the *TACO1*-KO A-P-E state it connected only to the P-site rRNA (Extended Data Fig. 9, a and b). Similarly, in the WT A-P state, nascent peptide density bridged both A- and P-site tRNAs, while in the *TACO1*-KO A-P state density extended to the P-site tRNA (Extended Data Fig. 9, c and d). Both A-P-E and A-P states in *TACO1*-KO cells also showed weaker density at the tip of the A-site tRNA compared to WT (Extended Data Fig. 9, b and d). These observations indicate that TACO1 stabilizes the A-site tRNA and promotes efficient peptidyl transfer.

Collectively, these findings suggest that TACO1 facilitates a critical step in mitochondrial translation elongation by competing with mtEF-Tu for mitoribosome binding and promoting the accommodation of the A-site tRNA. Through its interactions with the A-site tRNA and rRNA, TACO1 stabilizes the tRNA acceptor stem and promotes peptidyl transfer, thereby mediating an intermediate elongation step essential for resolving mitoribosome stalling at challenging translation motifs, such as polyproline stretches. Thus, TACO1 emerges as an important mediator that ensures both the efficiency of mitochondrial protein synthesis and the structural integrity of the 55S mitoribosome.

### Potential conserved mechanism of TACO1 family in translation

The TACO1 protein family, also known as the YebC family, is conserved and widely distributed across bacteria, archaea, and mitochondria (Extended Data Fig. 10a)^71^. In addition to human TACO1, bacterial orthologs such as YebC from *S. pyogenes* and YebC2 (formerly YeeI) from *B. subtilis* were recently identified as factors that relieve ribosome stalling^54,55^. To analyze whether bacterial orthologs operate through a conserved mechanism, we used AlphaFold^60^ to predict complex structures between *Sp*YebC and *Sp*bL12, as well as *Bs*YebC2 and *Bs*bL12. Both predicted complexes exhibited high confidence scores and displayed binding patterns closely resembling our experimentally determined human TACO1–bL12m interaction (Extended Data Fig. 10b), suggesting that bacterial TACO1 orthologs likely employ a conserved mechanism to promote elongation and resolve ribosome stalling.

Beyond translation, some TACO1 family members have also been implicated in transcriptional regulation via promoter-specific recognitions. Phylogenetic analysis of 731 identified TACO1 orthologs revealed three major clades (Extended Data Fig. 10a). Orthologs implicated in translation cluster within Clade III^23,38,54,55,72^; transcription-associated orthologs primarily populate Clade II^73–77^; and the first identified member ^71^, residing in Clade I, remains functionally uncharacterized. Conservation analysis shows that residues corresponding to those in TACO1 that contact the A-site tRNA acceptor stem—a double-helical element resembling double-stranded DNA—are highly conserved (Extended Data Fig. 11, a and d). By contrast, residues mediating interactions with 16S rRNA are more variable (Extended Data Fig. 11, b and c), possibly reflecting functional specialization within the family. Collectively, these findings suggest a conserved mechanistic role for some TACO1 family members in translation elongation, with other YebC-family members acting on transcriptional regulation potentially emerging in specific evolutionary branches.

## Discussion

Translation elongation is tightly regulated to ensure fidelity and efficiency, particularly when ribosomes encounter structurally or kinetically challenging sequences like polyproline tracts. These motifs slow peptide bond formation due to the unique rigidity and cyclic structure of proline, which hinders proper accommodation of Pro-tRNAs in both the P and A sites^25,33,78^. In bacteria and eukaryotes, elongation factors EF-P and eIF5A, respectively^24–26,30^, resolve this challenge by occupying the E-site and stabilizing the P-site tRNAs to promote transpeptidation^31–33^. Some ABCF proteins act similarly^34–37^ and can substitute for EF-P^27–29^, highlighting diverse cellular solutions to elongation stalls.

Mitochondria, however, lack EF-P, eIF5A, and ABCF orthologs, raising the question of how they navigate elongation challenges. Recent evidence identified TACO1 as a mitochondria-specific elongation factor that resolves stalling, particularly at polyproline-rich sequences such as the triple-proline stretch in CO1^23^. Despite its functional convergence with EF-P and related factors, TACO1 and its prokaryotic counterparts are structurally distinct, suggesting that compared to their bacterial ancestors, mitochondria have retained a single specialized strategy to overcome stalling during elongation.

Our high-resolution cryo-EM analysis reveals that TACO1 binds adjacent to the mitoribosomal A-site, establishing extensive interactions with the 16S rRNA and bL12m of the mtLSU, the acceptor stem of the A-site tRNA, and bS12m on the mtSSU. This unique binding site partially overlaps with that of mtEF-Tu. In the absence of TACO1, we observed a striking accumulation of mitoribosomes in A/T states with persistent mtEF-Tu binding, and elevated free subunits. These observations indicate that the A/T-tRNA fails to efficiently transit and/or stably accommodate into the A-site, thereby impairing transpeptidation, stalling elongation and triggering mitoribosome subunit dissociation. We propose that TACO1 acts as a translation elongation regulator that resolves mitoribosomal stalling by competing with mtEF-Tu for mitoribosome binding which may influence mtEF-Tu release, stabilizing the A-site tRNA accommodation, and enhancing peptidyl transfer efficiency to facilitate elongation resumption (Fig. 4).

**Fig. 4.**
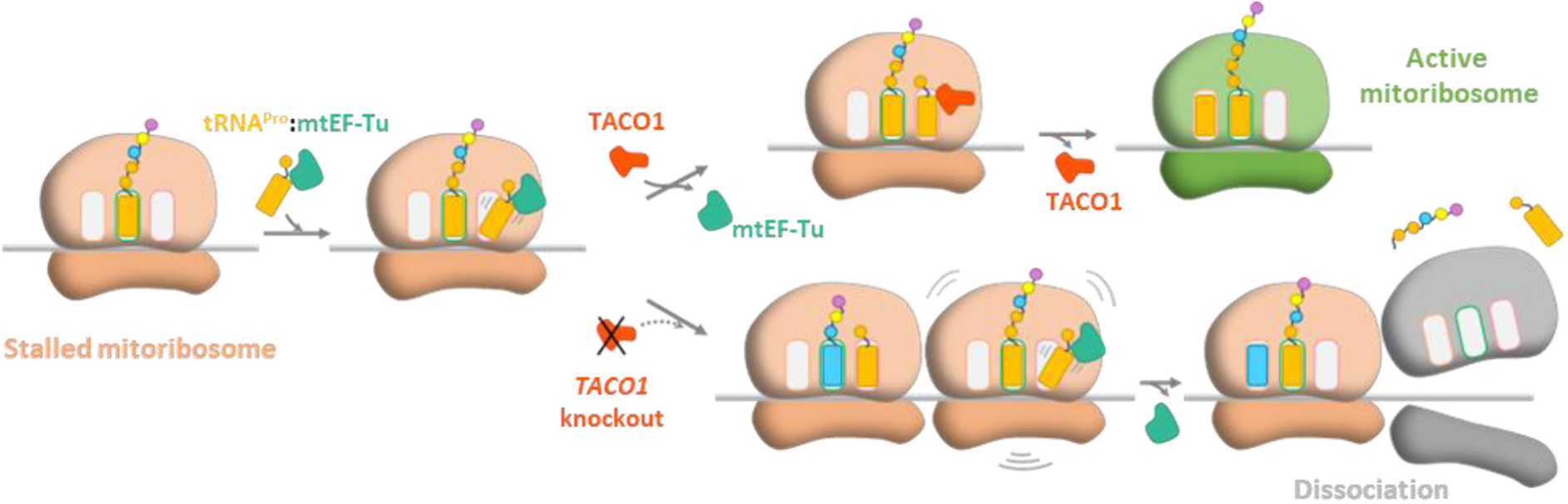
| Proposed Mechanism of TACO1 in alleviating translation stalling. Translation stalling occurs when mtEF-Tu-bound tRNAs fail to efficiently transition from the A/T site to the A site, often due to specific nascent peptide features such as polyproline motifs. TACO1 resolves this stalling by competing with EF-Tu for mitoribosome binding, thereby promoting mtEF-Tu release, stabilizing the acceptor stem of the A-site tRNA, and facilitating peptidyl transfer to restore translational elongation. In the absence of TACO1 (*TACO1*-KO cells), prolonged mtEF-Tu residence in A/T states, persistent stalling at challenging motifs leads to mitoribosomal subunit dissociation and the release of truncated polypeptides.

Comparative biochemical analyses of mitochondrial and *E. coli* translation elongation highlight fundamental adaptations in the mammalian system. In bacteria, EF-Tu·GTP tightly binds aminoacyl-tRNA to form a highly stable ternary complex that delivers the aa-tRNA to the ribosomal A-site, followed by rapid GTP hydrolysis and GDP–GTP exchange via EF-Ts. In contrast, mammalian mtEF-Tu exhibits ∼2 orders of magnitude weaker nucleotide affinity (K_d_ ≈ 19 ± 9 µM)^66^ but retains nanomolar stability for ternary complex formation with mitochondrial aa-tRNAs (Kd of ∼18 ± 4 nM)^79^. This shift in kinetic parameters makes guanine nucleotide exchange rate-limiting, underscoring the role of mtEF-Ts in maintaining elongation efficiency. In this context, the 25 µM GTP affinity measured for TACO1, comparable to that of mtEF-Tu, suggests physiologically relevant GTP binding. However, it does not by itself imply GTPase activity, which we did not detect robustly *in vitro*. Within the specialized mitochondrial framework, our data suggest that TACO1 could act as an auxiliary component of the elongation cycle rather than a canonical GTPase. Four non-mutually exclusive models can be proposed: (i) mtEF-Tu/mtEF-Ts tuner or scaffold: by interacting with mtEF-Ts, TACO1 could stabilize or accelerate GDP-GTP exchange on mtEF-Tu, facilitating efficient recharging of the elongation factor. (ii) Ribosome-proximal accelerator: TACO1 may occupy a site adjacent to mtEF-Tu on the mitoribosome, promoting rapid aa-tRNA accommodation or mtEF-Tu turnover under slow-decoding conditions. (iii) Anti-stall factor: TACO1 could reduce ribosomal pausing at problematic sequences (*e.g*., poly-Pro stretches) by favoring mtEF-Tu rebinding or engagement of the translocation factor mtEF-G1, thereby accelerating elongation without catalyzing GTP hydrolysis. (iv) Functional bridge between mtEF-Tu and mtEF-G modules: TACO1 might coordinate the transition from aa-tRNA delivery to translocation, aligning factor exchange dynamics with the kinetic landscape of mitochondrial translation. Discriminating among these models warrant future investigations. Together, they position TACO1 as a mitochondrial elongation modulator-potentially integrating GTP sensing, mtEF-Tu cycling, and ribosome dynamics to fine-tune translation efficiency within the constrained energetic environment of the organelle. Furthermore, large excess of TACO1 disrupts mitochondrial translation^22^, likely by obstructing the tRNA entrance channel, suggesting that timely dissociation of TACO1 and the stoichiometry of translation elongation factors are critical for avoiding elongation gridlock and maintaining translational throughput. Identifying the triggers and timing of TACO1 dissociation remains a key challenge for understanding how elongation is dynamically regulated within mitochondria.

TACO1 orthologs are widespread in mitochondria, bacteria, and archaea, but absent from the eukaryotic cytoplasm. Notably, in eukaryotes that have lost mitochondrial translation, such as *Plasmodium falciparum*, TACO1 orthologs are also missing, reinforcing its specialized role in mitoribosomal translation. In bacteria like *Streptococcus pyogenes*^54^ and *Bacillus subtilis*^55^, TACO1 orthologs—YebC/YebC2—similarly resolve ribosome stalling. Some bacterial species encode multiple paralogs with evidence of functional divergence^55^. For instance, in *E. coli*, *Ec*YebC binds to gene promoters and has been implicated in transcriptional regulation^76^, whereas *Ec*YeeN, closely related to *Bs*YebC2 phylogenetically (Extended Data Fig. 10a), may function in translation. Similar diversification is seen in *B. subtilis* and likely other bacterial species^55^. Our work lays the foundation for dissecting the molecular determinants of functional divergence within the TACO1 protein family and for exploring their potentials in health and disease, as antibiotic targets to inhibit virulent bacteria growth or as agonists to regulate mitochondrial translation.

The discovery of TACO1 as a dedicated elongation regulator in mitochondria not only reveals an ancient solution to the universal problem of polyproline-mediated stalling but also illustrates how translation systems independently evolve and adapt specialized machineries to preserve elongation under stress. In this light, TACO1 emerges as a mitoribosome-specific elongation factor, one that operates outside the canonical E-site paradigm to ensure the continuity of protein synthesis in the most challenging sequence contexts. Future work dissecting the regulation, dynamics, and evolutionary diversity of TACO1 will provide a critical window into how elongation is customized for the unique demands of mitochondrial translation.

## Acknowledgements

We thank V. Singh, A. Khawaja, and J. Rorbach at Karolinka Institutet for helpful discussions and providing purified mtEF-Tu. We thank J. Lin, K. Zhou, and S. Wu of the Yale Cryo-EM Resource for expert training and assistance with grid screening and data collection. We are grateful to N. Grigorieff and J. Elferich from University of Massachusetts for sharing the latest version of cisTEM. We also thank all members of the Xiong laboratory for assistance and encouragement throughout this project. Special thanks to S. Tang, A. Didychuk, I. Lomakin, and J. Wang for insightful discussions and suggestions. This work was supported by startup funds from Yale University to Y. Xiong, National Institute of General Medicine (NIGMS) grant R35-GM118141 to A. Barrientos, Muscular Dystrophy Association Research grant 1069392 to A. Barrientos, Muscular Dystrophy Association Development grant 22-1288334 to M. Brischigliaro and AFM-Téléthon Trampoline grant 28651 to M. Brischigliaro.

## Author contributions

Y.X. and A.B. conceived and supervised this study. S.W. performed cell culture and mitochondria isolation. S.W., C.W. and W.Z. carried out cryo-EM grid screening and data collection. S.W. and Y.X. processed the cryo-EM data. S.W. and Y.Z. identified the TACO1 and built the structural models. M.B. performed the GTP-binding assays, sucrose gradient sedimentation analyses and mass spectrometry experiments. S.W., Y.X., M.B. and A.B. contributed to drafting the manuscript. All authors participated in data analysis and manuscript review.

## Data and materials availability

The cryo-EM density maps of the mitoribosome–TACO1 complex generated in this study have been deposited in the Electron Microscopy Data Bank under accession numbers: EMD-70621 (consensus map of WT 55S mitoribosome), EMD-70620 (focused mtSSU map), EMD-70619 (focused mtLSU map), and EMD-70592 (composite map of the WT 55S mitoribosome). The corresponding molecular model has been deposited in the Protein Data Bank under accession number PDB 9OLF (55S mitoribosome in complex with LRPPRC, SLIRP, OXA1L, and TACO1). Mass spectrometry data have been deposited in the PRIDE database under accession PXD064026 (Reviewers can access the dataset by logging in to the PRIDE website using Project accession: PXD064026, Token: oTJOZvo4r8cQ, or Username: Password: GiLD0fQdMAXP). Source data and other materials are available from the authors upon reasonable request.

---

**Extended Data Fig. 1.**
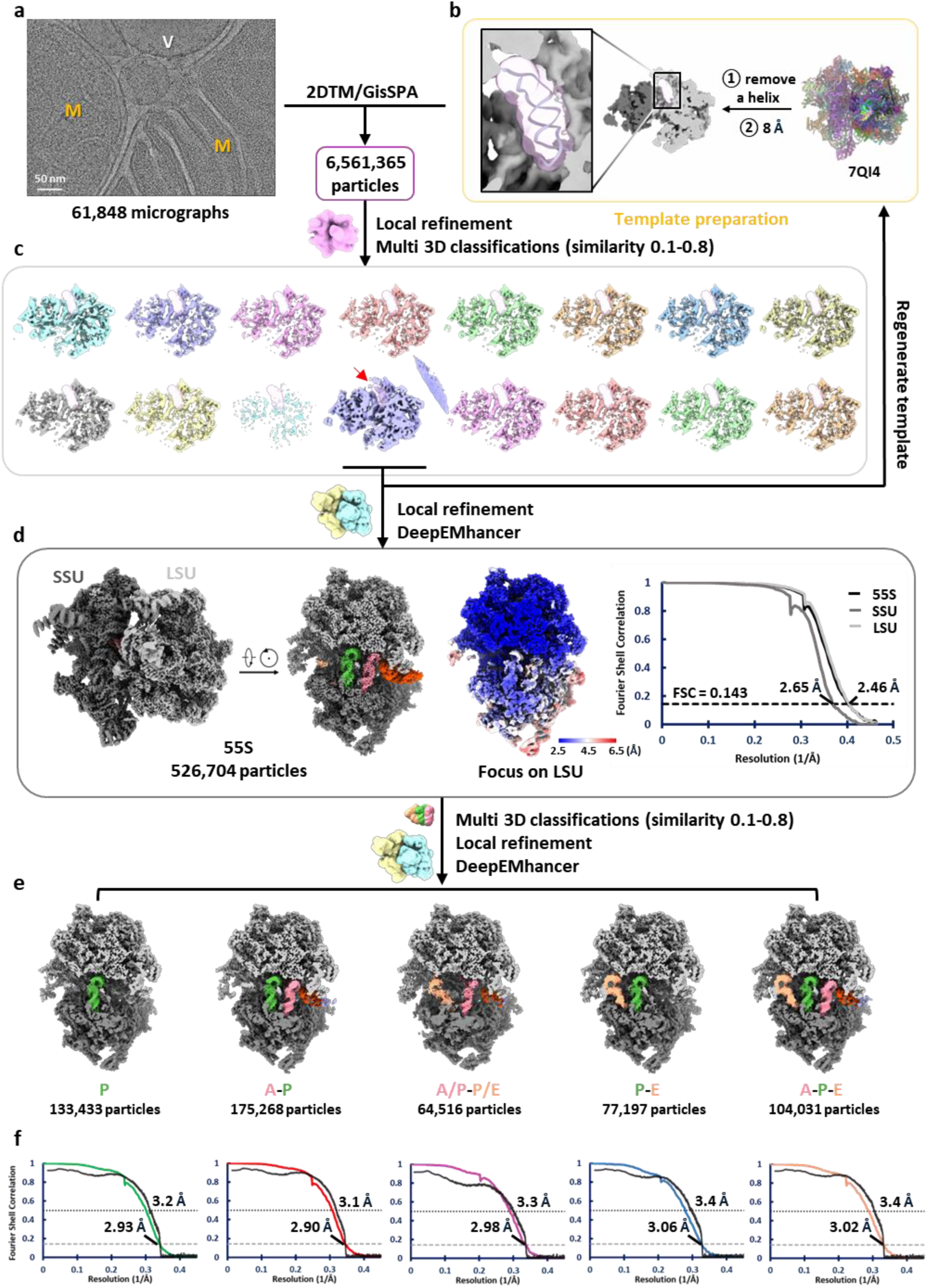
| Workflow of single-particle cryo-EM analysis of the human mitoribosome within mitochondria. Particles were initially picked from raw micrographs (**a**) via 2DTM and GisSPA guided by a low-resolution template (**b**), which was generated using PDB ID: 7QI4 with a helical region of the 16S rRNA removed. (**c**) Following 3D classification, the class, in which the removed helix was restored (read arrow), was selected to create an updated low-resolution template, again omitting the helix, for integrative particle re-picking. (**d**) Particles from all classes showing helix restoration were combined to reconstruct the consensus cryo-EM map. (**e**) Different mitoribosomal states were resolved through focused 3D classification on the tRNA region. (**f**) Fourier shell correlation (FSC) curves for the final reconstructions (colored). Model vs. map FSCs are shown in black.

**Extended Data Fig. 2.**
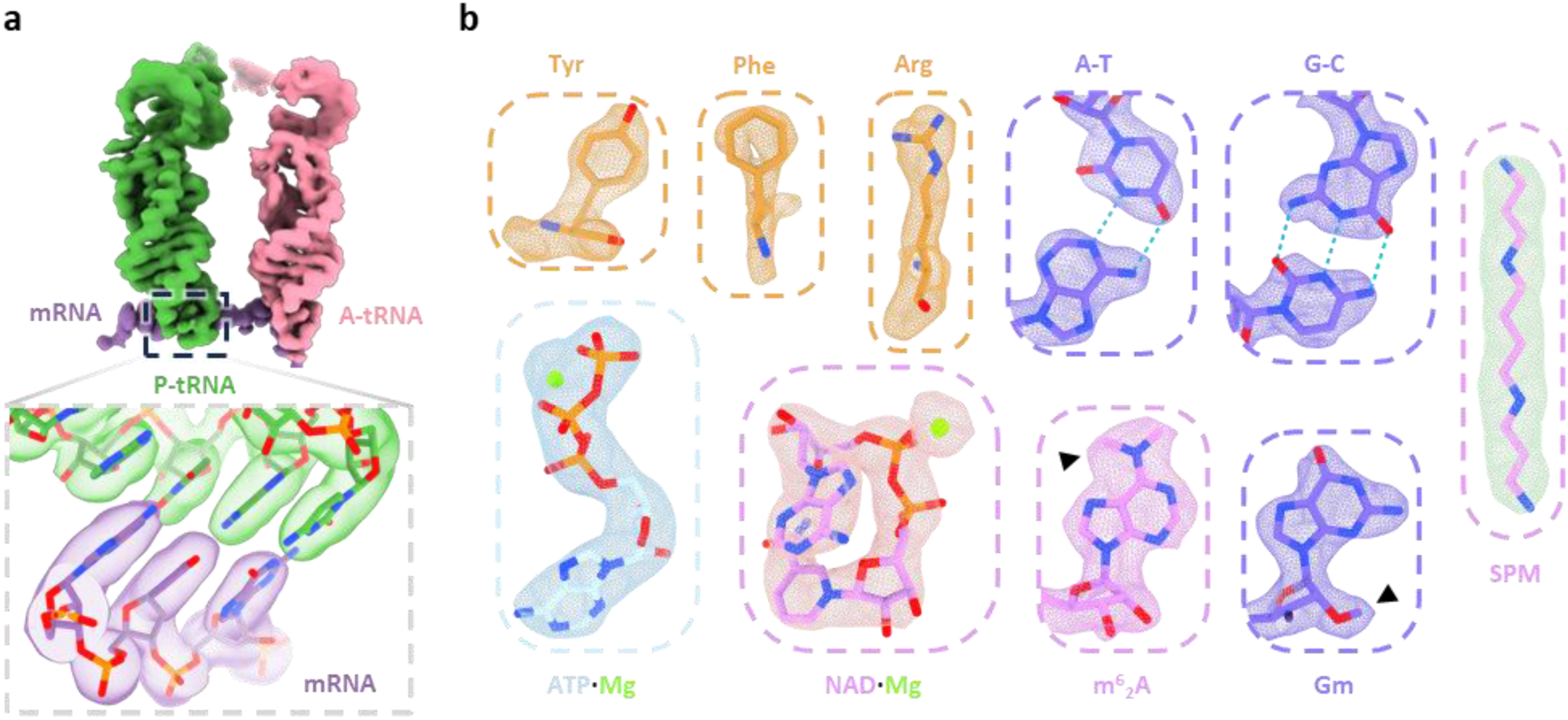
| High-resolution structural features of the *in organello* human mitoribosome consensus cryo-EM map. (**a**) Densities of the tRNAs and mRNA, with an enlarged view highlighting the codon–anticodon pairing between the mRNA and the P-site tRNA. (**b**) Representative high-resolution structural details, including resolved amino acid side chains, RNA base pairs, base modifications, and bound endogenous ligands. Modifications and ligands are labeled: Gm, 2’-O-methylguanosine; m^62^A, N^6^,N^6^-dimethyladenosine; SPM, spermine.

**Extended Data Fig. 3.**
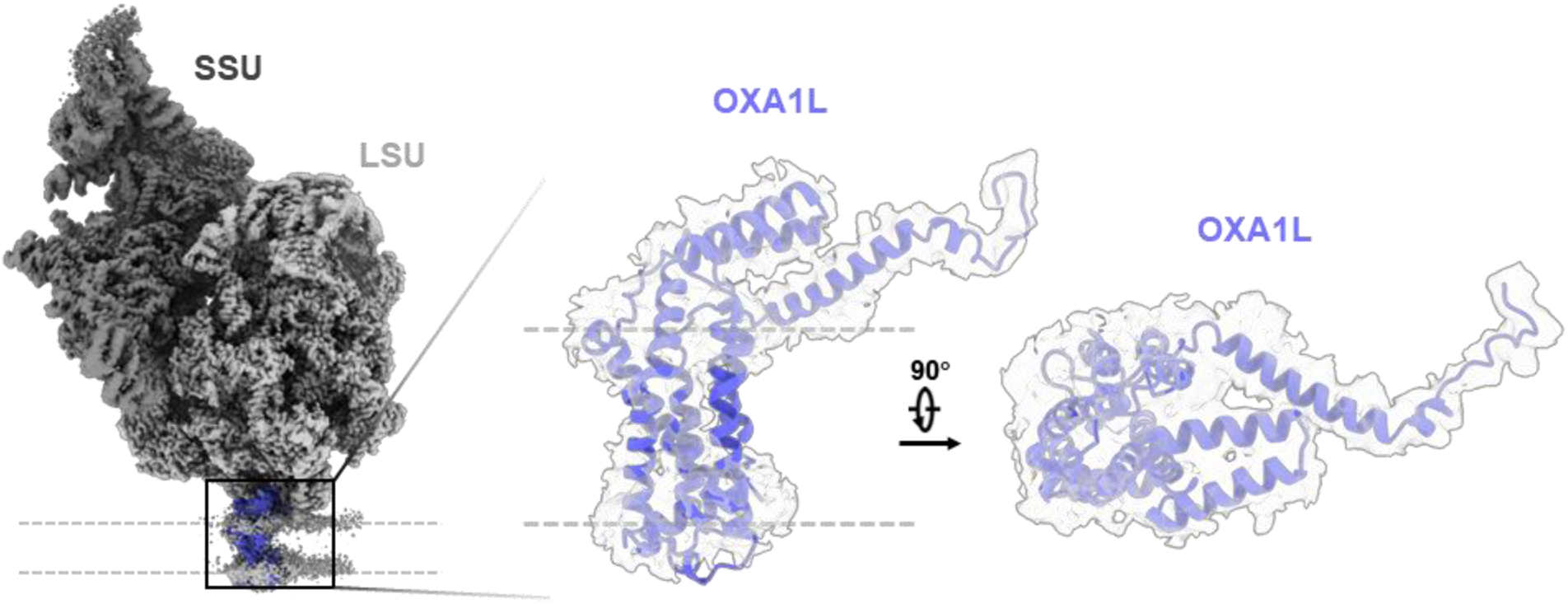
| OXA1L density at the mitoribosome–membrane interface. Consensus cryo-EM map of the mitoribosome bound to the inner mitochondrial membrane, with mtSSU (dark grey), mtLSU (light grey), and additional density at the end of the polypeptide exit tunnel fitted with the AlphaFold-predicted OXA1L model (blue). Side and top views highlight the fit of the OXA1L model into the density.

**Extended Data Fig. 4.**
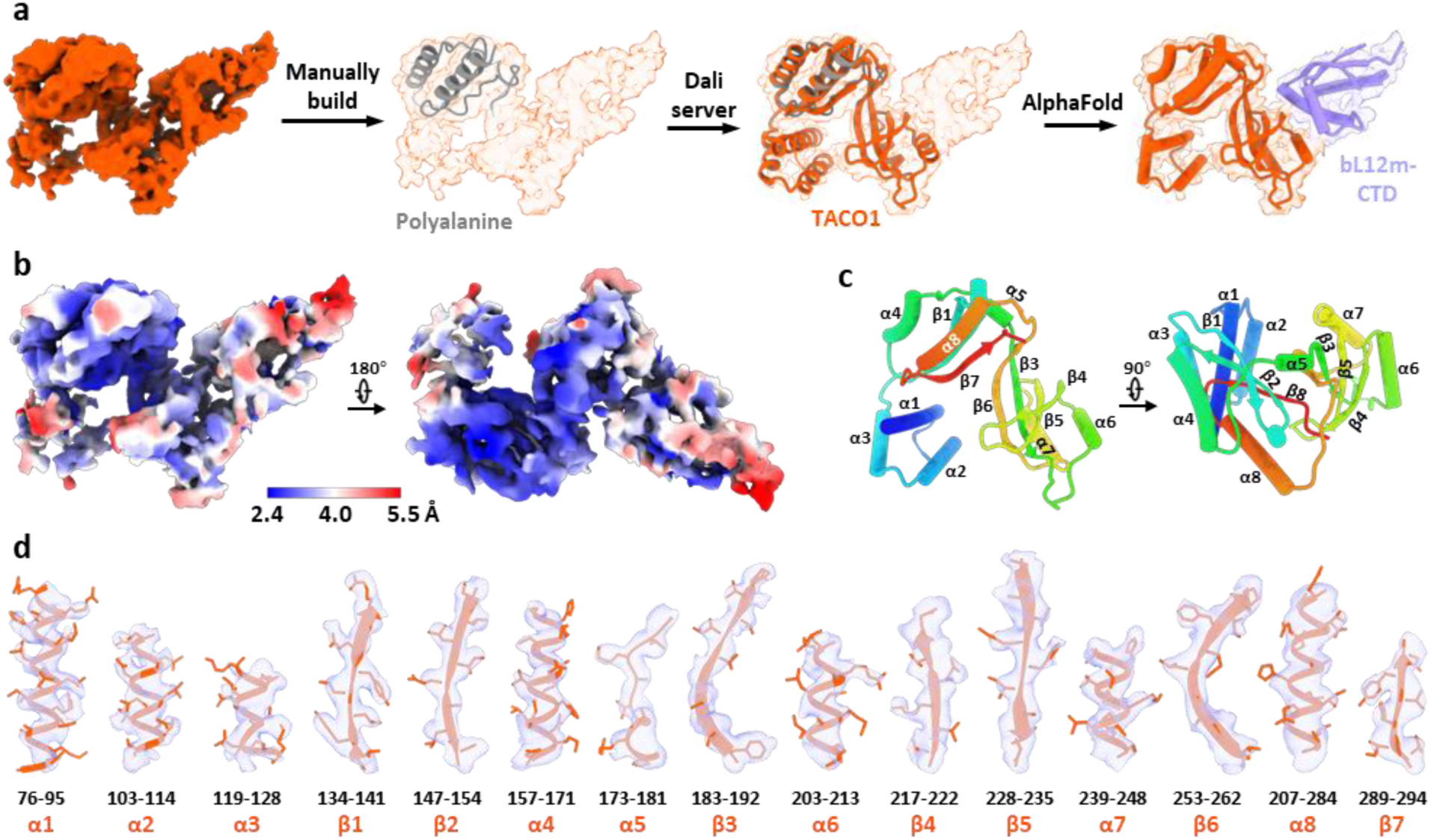
| Identification of TACO1 complex with the C-terminal domain of bL12m. (**a**) Workflow for identifying the extra density as TACO1 (orange) and C-terminal domain of bL12m (bL12m-CTD) (light blue). (**b**) Local resolution map of the extra density, highlighting the quality of the reconstruction. (**c**) Assigned secondary structural elements of TACO1. (**d**) Close-up view showing the fit of all TACO1 structural elements into the cryo-EM density.

**Extended Data Fig. 5.**
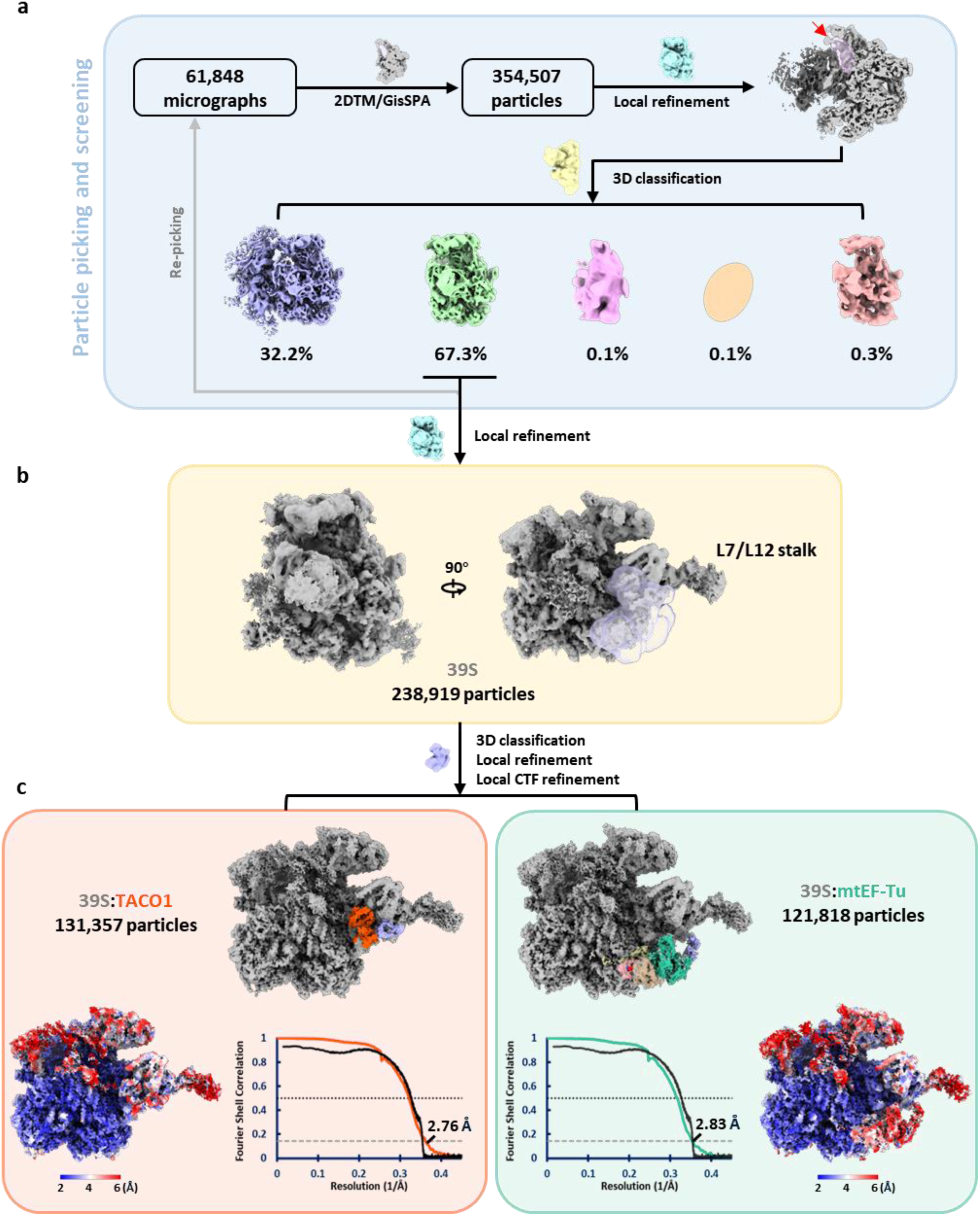
| Data processing workflow for free 39S mitoribosomes (mtLSU) from wide-type human cells. (**a**) Particles were picked using a mitoribosome template lacking a 16S rRNA helix, selecting for particles that restored the helix but lacked the SSU. (**b**) Extra density detected near the A-site by the L7/L12 stalk. (**c**) 3D classification focused on the extra density resolved two distinct 39S mitoribosome classes, one associated with TACO1 (left) and the other with mtEF-Tu (right). Bottom panels show local resolution maps and Fourier shell correlation (FSC) curves for each class.

**Extended Data Fig. 6.**
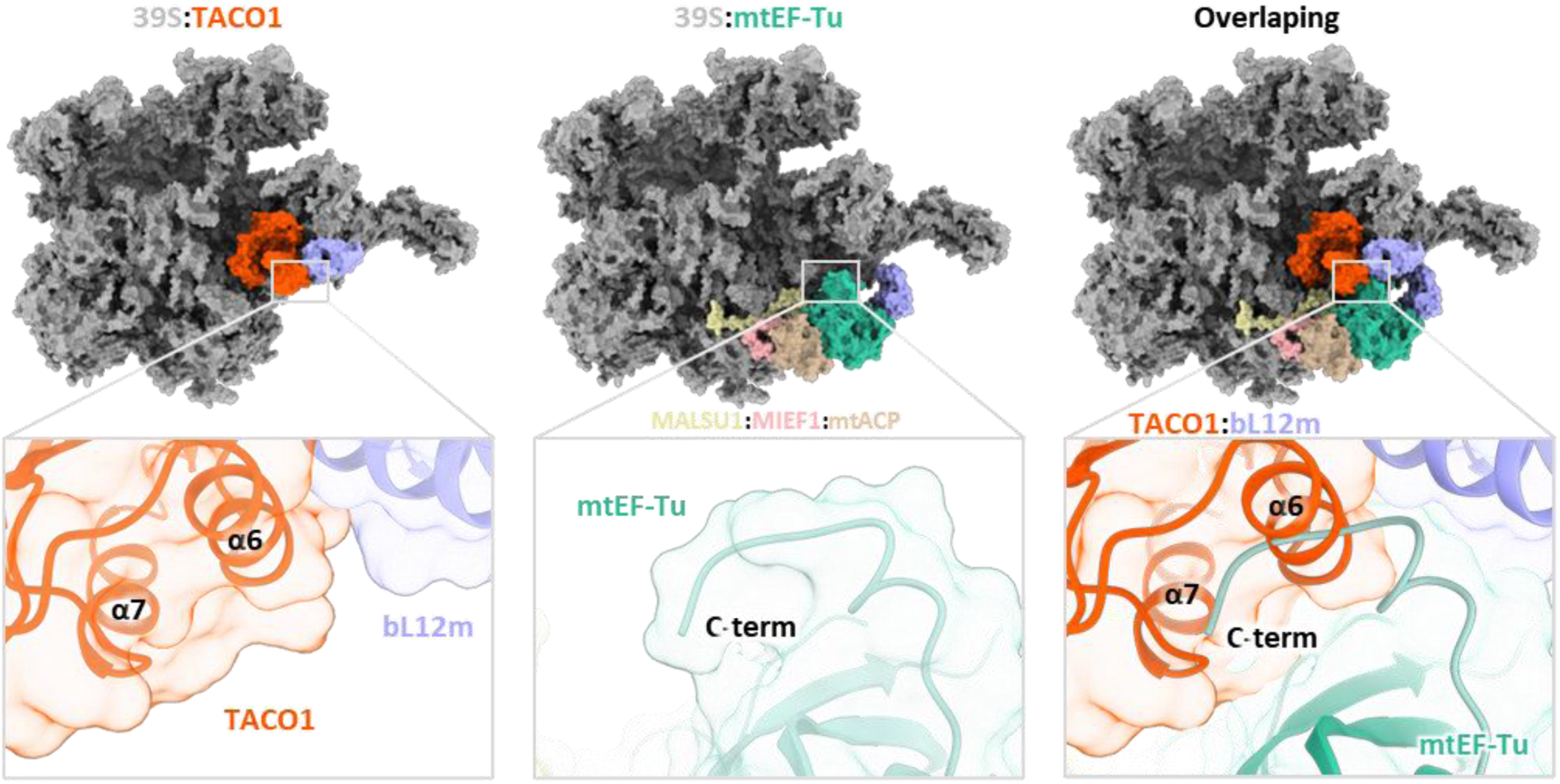
| Structural comparison of TACO1 and mtEF-Tu binding sites on the 39S mitoribosome. Superposition of the TACO1-bound (left) and the mtEF-Tu-bound (middle) 39S mitoribosome structures reveals steric clashes between the two proteins (right), indicating that their binding to the 39S mitoribosome is mutually exclusive.

**Extended Data Fig. 7.**
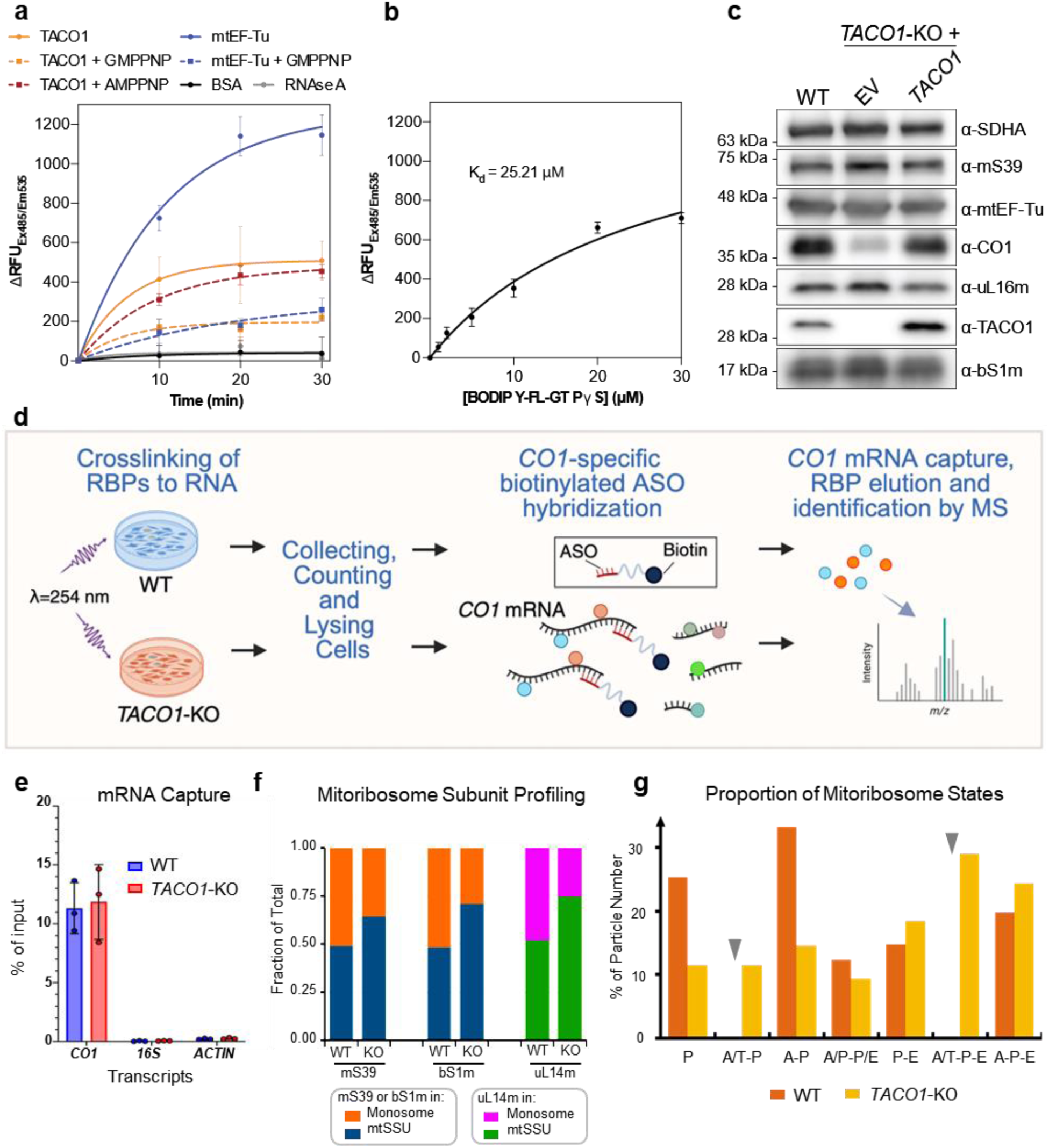
| GTP binding properties of TACO1 and its impact on *CO1* mRNA-binding proteins and mitoribosome association in *TACO1*-KO cells. (**a**) BODIPY-FL-GTPγS binding assay of recombinant mtEF-Tu and TACO1, with or without preincubation with GMPPNP or AMPPNP (*n* = 3). (**b**) Guanine nucleotide-binding affinity to TACO1 measured using increasing concentrations of BODIPY-FL-GTPγS (0 μΜ, 1 μΜ, 2 μΜ, 5 μΜ, 10 μΜ, 20 μΜ and 30 μΜ) incubated with 500 nM TACO1 for 10 min. Data represented as average of triplicates (n=3) with error bars (SD). The dissociation constant (K_D_) was determined by fitting nonlinear curve regression using GraphPad Prism 10. (**c**) Steady-state levels of mitoribosome subunits and mtEF-Tu in WT and *TACO1*-KO, and rescued KO cells. **(d)** Workflow for the UV-induced protein-RNA crosslinking, *CO1* mRNA pulldown and quantitative LC-MS/MS analysis of *CO1* mRNA-specific binding proteins comparing TACO1-KO and WT cells. (**e**) Specificity of the approach assessed by qPCR quantification of *CO1* mRNA relative to mitochondrial 16S rRNA or ACTIN transcripts. Data are plotted as mean ± SD (*n* = 3). (**f**) Densitometric quantitative analysis of the distribution of each mitoribosome marker in Fig. 3d across the sedimentation fractions corresponding to the dissociated subunits (F10 + F11 for the mtSSU, and F8 + F9 for the mtLSU) and monosomes (F5 + F6), presented as the fraction of protein in each structure. (**g**) Particle distribution of mitoribosomes across various translation elongation states in WT and *TACO1*-KO cells. Grey arrows mark states for which no mitoribosomes were identified.

**Extended Data Fig. 8.**
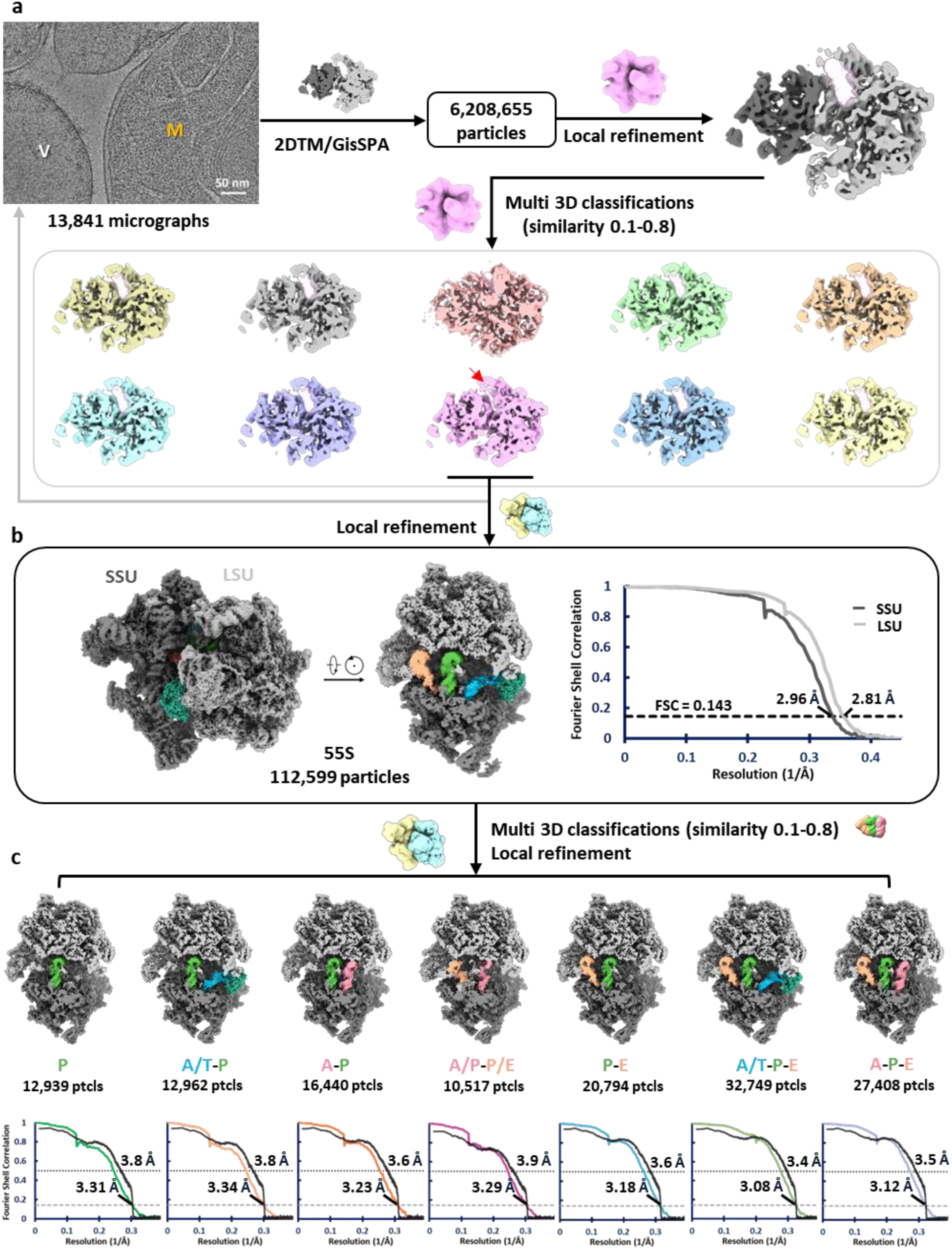
| Cryo-EM single-particle analysis workflow for the human mitoribosome from *TACO1*-KO cells. (**a**) Template-based particle picking followed by iterative refinement through 3D classification. (**b**) Composite map of the 55S mitoribosome reconstructed by focused refinements on the mtLSU or mtSSU separately, along with the corresponding Fourier shell correlation (FSC) curves. (**c**) Structural states of the mitoribosome identified in *TACO1*-KO cells and their associated FSC curves.

**Extended Data Fig. 9.**
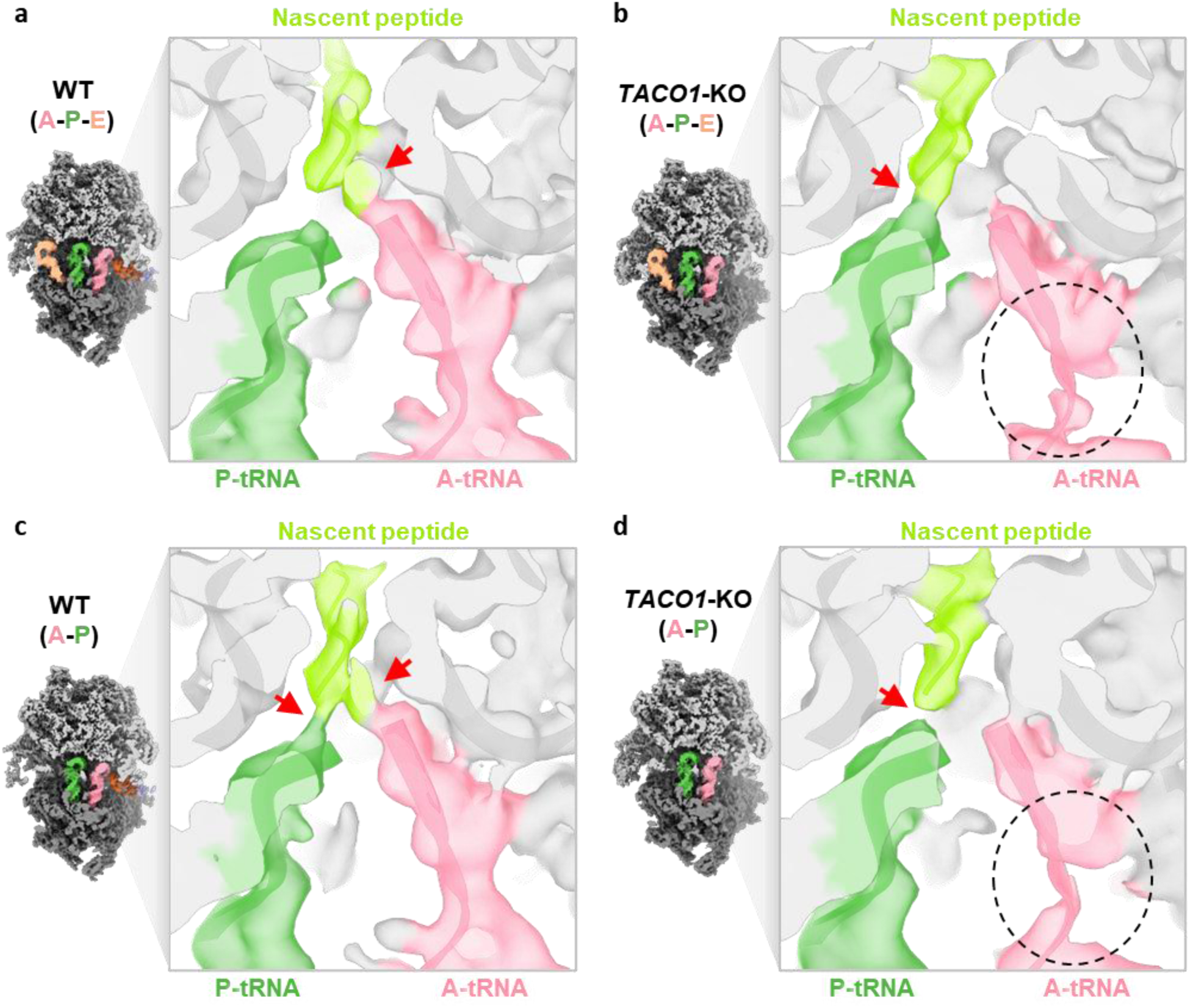
| Density differences at the peptidyl transfer center in WT and *TACO1*-KO mitoribosomes. Cryo-EM maps of the A-P-E state (**a**, **b**) and A-P state (**c**, **d**) from WT cells and *TACO1*-KO cells, displayed at a 3σ contour. Densities are colored according to the fitted models: A-site tRNA (pink), P-site tRNA (green), and nascent peptide (yellow-green). Red arrows mark connections between tRNAs and the nascent peptide, and black circles indicate flexible regions with weaker density.

**Extended Data Fig. 10.**
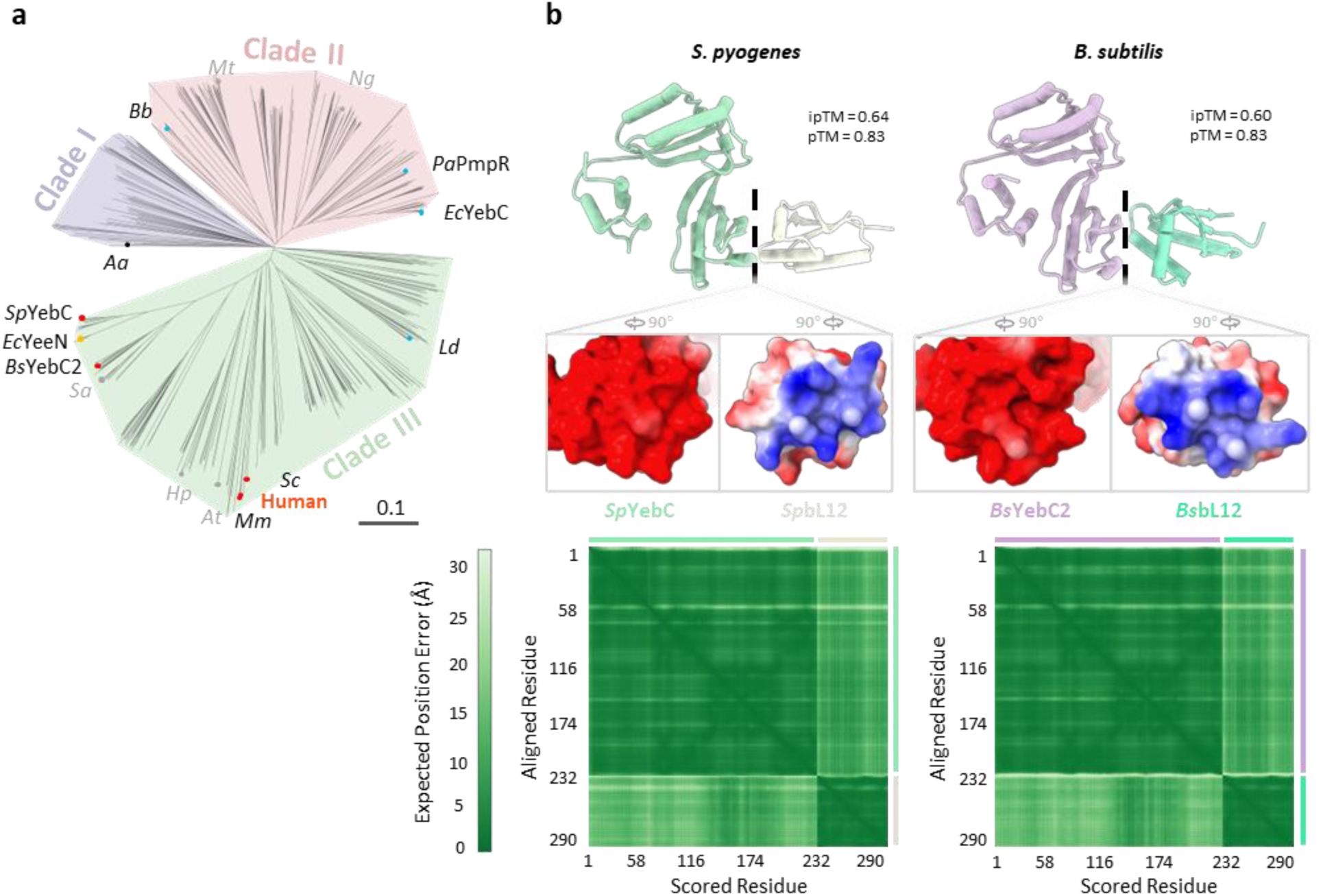
| Phylogenetic and structural analysis of TACO1 orthologs across diverse species. (**a**) Phylogenetic tree of 731 TACO1 orthologs classified into three major clades (83/288/360). Branches corresponding to research model organisms are marked with dots. Red dots indicate orthologs implicated in translation, and blue dots indicate those associated with transcriptional regulation. Species abbreviations: *Mm*, *Mus musculus* (mouse); *Sc*, *Saccharomyces cerevisiae* (yeast); *At*, *Arabidopsis thaliana*; *Hp*, *Helicobacter pylori*; *Sa*, *Staphylococcus aureus*; *Ec*, *Escherichia coli*; *Sp*, *Streptococcus pyogenes*; *Bs*, *Bacillus subtilis*; *Ld*, *Lactobacillus delbrueckii*; *Aa*, *Aquifex aeolicus*; *Bb*, *Borreliella burgdorferi*; *Mt*, *Mycobacterium tuberculosis*; *Ng*, *Neisseria gonorrhoeae*; *Pa*, *Pseudomonas aeruginosa*. (**b**) Predicted complex structures (top) of TACO1 and bL12 orthologs from *S. pyogenes* and *B. subtilis*, along with their corresponding electrostatic surface potentials at the interaction interfaces (middle) and the predicted aligned error matrices (bottom).

**Extended Data Fig. 11.**
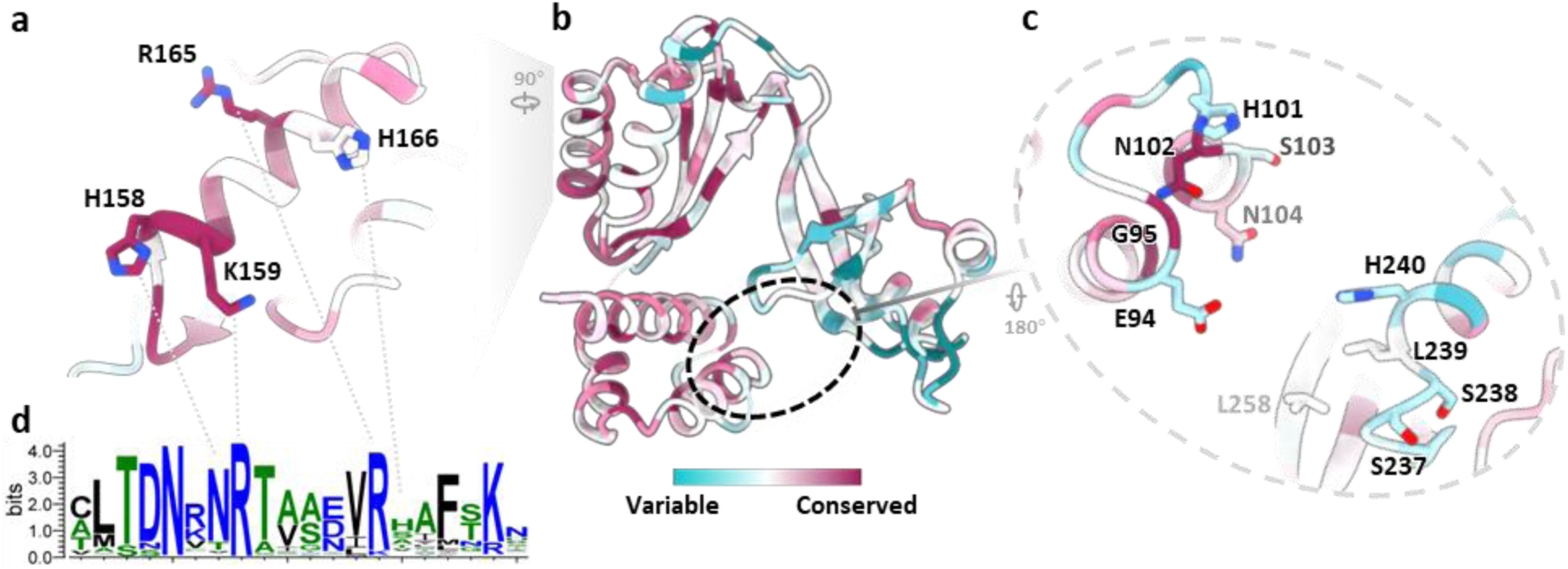
| Conservation analysis of TACO1 orthologs. (**a-c**) Overall structure of hTACO1 colored according to evolutionary conservation scores across 731 TACO1 orthologs (**b**), with close-up views of the conserved residues at the interface with tRNA (**a**) and 16S rRNA (**c**). (**d**) Sequence logos depicting sequence conservation (in bits) derived from a multiple sequence alignment of the 731 TACO1 orthologs.

**Extended Data Table 1.** | Data collection and refinement statistics.

|  | 55S mitoribosome from WT |  |  |  |
| --- | --- | --- | --- | --- |
|  | Consensus<br>(EMD-70592)<br>(PDB 9OLF) | P<br>(EMD-71630)<br>(PDB 9PGF) | A-P<br>(EMD-71634)<br>(PDB 9PGL) | A/P-P/E<br>(EMD-71829)<br>(PDB 9PSM) |
| **Data collection and processing** |  |  |  |  |
| Magnification | 81,000 | 81,000 | 81,000 | 81,000 |
| Voltage (kV) | 300 | 300 | 300 | 300 |
| Electron exposure (e-/Å<sup>2</sup>) | 50 | 50 | 50 | 50 |
| Defocus range (µm) | 1.8-2.0 | 1.8-2.0 | 1.8-2.0 | 1.8-2.0 |
| Pixel size (Å) | 1.068 | 1.068 | 1.068 | 1.068 |
| Final Micrographs (no.) | 61,848 | 61,848 | 61,848 | 61,848 |
| Initial particle images (no.) | 6,561,365 | 6,561,365 | 6,561,365 | 6,561,365 |
| Final particle images (no.) | 526,704 | 133,433 | 175,268 | 64,516 |
| Symmetry imposed | C1 | C1 | C1 | C1 |
| Map resolution (Å) | 2.46 | 2.93 | 2.9 | 2.98 |
| FSC threshold | 0.143 | 0.143 | 0.143 | 0.143 |
| **Refinement** |  |  |  |  |
| Model resolution (Å) | 2.6 | 3.2 | 3.1 | 3.2 |
| FSC threshold | 0.5 | 0.5 | 0.5 | 0.5 |
| Model composition |  |  |  |  |
| Non-hydrogen atoms | 192,936 | 178,740 | 180,184 | 181,859 |
| Protein residues | 16,295 | 14,888 | 14,888 | 15,103 |
| Nucleotide | 2,828 | 2,690 | 2,758 | 2,757 |
| Ligands | 277 | 276 | 277 | 277 |
| *B* factors (Å<sup>2</sup>) |  |  |  |  |
| Protein | 117 | 168 | 150. | 157 |
| Nucleotide | 90 | 140 | 126 | 133 |
| Ligand | 68 | 119 | 105 | 106 |
| R.m.s. deviations |  |  |  |  |
| Bond lengths (Å) | 0.005 | 0.003 | 0.006 | 0.005 |
| Bond angles (°) | 0.8 | 0.5 | 0.8 | 0.8 |
| Validation |  |  |  |  |
| MolProbity score | 1.1 | 1.2 | 1.2 | 1.3 |
| Clashscore | 2.9 | 3.5 | 3.8 | 4.2 |
| Poor rotamers (%) | 0.01 | 0.01 | 0.01 | 0.01 |
| Ramachandran plot |  |  |  |  |
| Favored (%) | 97.84 | 97.85 | 97.67 | 97.42 |
| Allowed (%) | 2.13 | 2.14 | 2.32 | 2.58 |
| Disallowed (%) | 0.04 | 0.01 | 0.01 | 0.01 |

**Extended Data Table 2.** | Data collection and refinement statistics.

|  | 55S mitoribosome from WT |  | 39S mitoribosome from WT |  |
| --- | --- | --- | --- | --- |
|  | P-E | A-P-E | 39S:TACO1 | 39S:mtEF-Tu |
|  | (EMD-71623) | (EMD-71633) | (EMD-71797) | (EMD-71802) |
|  | (PDB 9PG8) | (PDB 9PGI) | (PDB 9PR4) | (9PRA) |
| **Data collection and processing** |  |  |  |  |
| Magnification | 81,000 | 81,000 | 81,000 | 81,000 |
| Voltage (kV) | 300 | 300 | 300 | 300 |
| Electron exposure (e-/Å<sup>2</sup>) | 50 | 50 | 50 | 50 |
| Defocus range (µm) | 1.8-2.0 | 1.8-2.0 | 1.8-2.0 | 1.8-2.0 |
| Pixel size (Å) | 1.068 | 1.068 | 1.068 | 1.068 |
| Final Micrographs (no.) | 61,848 | 61,848 | 61,848 | 61,848 |
| Initial particle images (no.) | 6,561,365 | 6,561,365 | 6,561,365 | 6,561,365 |
| Final particle images (no.) | 77,197 | 104,031 | 131,357 | 121,818 |
| Symmetry imposed | C1 | C1 | C1 | C1 |
| Map resolution (Å) | 3.06 | 3.02 | 2.77 | 2.83 |
| FSC threshold | 0.143 | 0.143 | 0.143 | 0.143 |
| **Refinement** |  |  |  |  |
| Model resolution (Å) | 3.4 | 3.4 | 3.0 | 3.1 |
| FSC threshold | 0.5 | 0.5 | 0.5 | 0.5 |
| **Model composition** |  |  |  |  |
| Non-hydrogen atoms | 182,250 | 183,684 | 111,602 | 115,757 |
| Protein residues | 15,140 | 15,140 | 9,453 | 9,968 |
| Nucleotide | 2,760 | 2,828 | 1,630 | 1,630 |
| Ligands | 276 | 276 | 183 | 184 |
| **B factors (Å<sup>2</sup>)** |  |  |  |  |
| Protein | 195 | 167 | 131 | 148 |
| Nucleotide | 168 | 141 | 110 | 118 |
| Ligand | 144 | 114 | 90 | 112 |
| **R.m.s. deviations** |  |  |  |  |
| Bond lengths (Å) | 0.003 | 0.008 | 0.005 | 0.004 |
| Bond angles (°) | 0.5 | 0.9 | 0.8 | 0.8 |
| **Validation** |  |  |  |  |
| MolProbity score | 1.3 | 1.3 | 1.0 | 1.1 |
| Clashscore | 4.2 | 3.6 | 1.9 | 2.6 |
| Poor rotamers (%) | 0.00 | 0.01 | 0.00 | 0.02 |
| **Ramachandran plot** |  |  |  |  |
| Favored (%) | 97.60 | 97.30 | 97.87 | 97.69 |
| Allowed (%) | 2.40 | 2.68 | 2.12 | 2.30 |
| Disallowed (%) | 0.01 | 0.02 | 0.01 | 0.01 |

**Extended Data Table 3.**
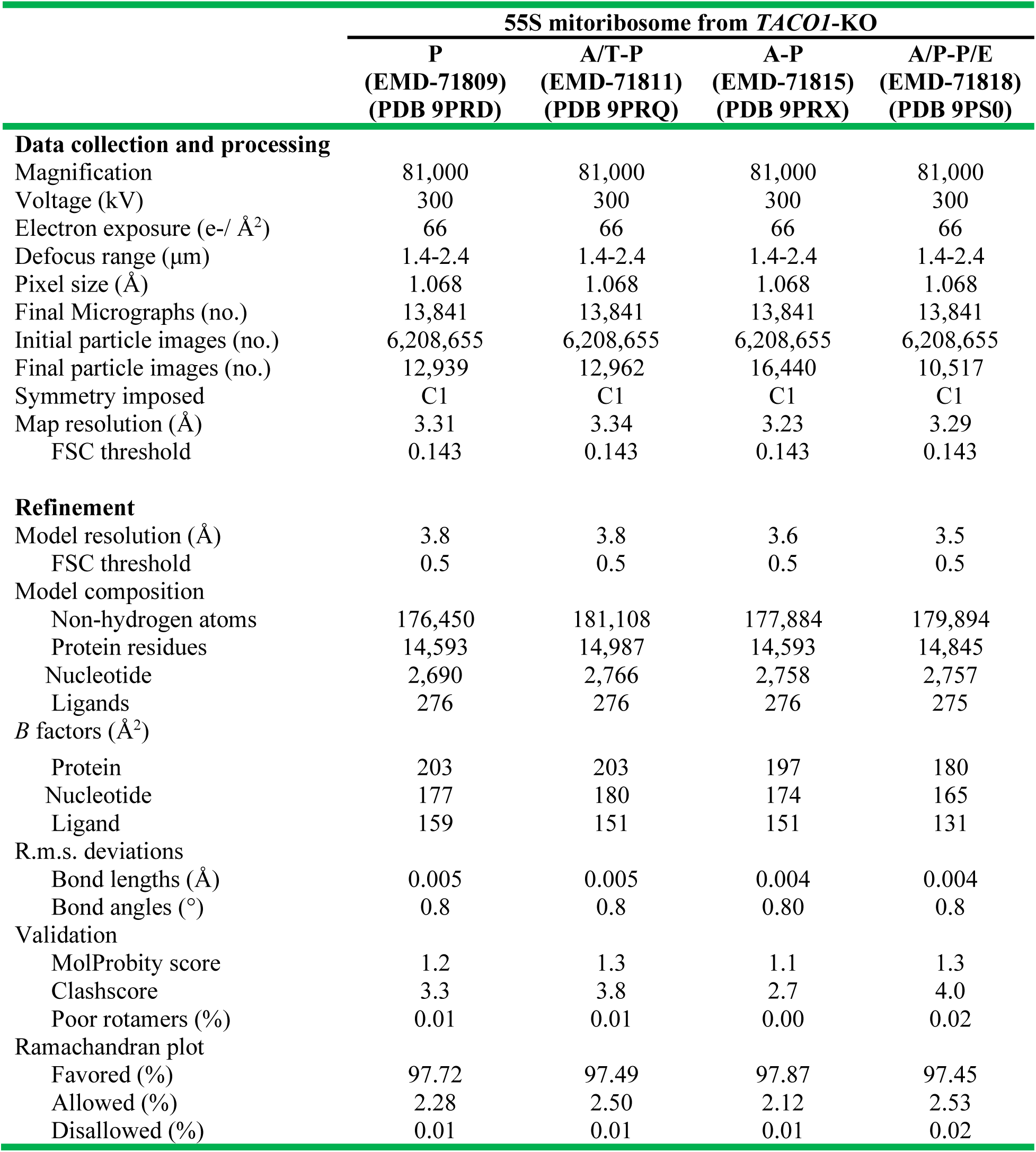
| Data collection and refinement statistics.

**Extended Data Table 4.** | Data collection and refinement statistics.

|  | 55S mitoribosome from *TACO1*-KO |  |  |
| --- | --- | --- | --- |
|  | P-E<br>(EMD-71635)<br>(PDB 9PGM) | A/T-P-E<br>(EMD-71825)<br>(PDB 9PS7) | A-P-E<br>(EMD-71828)<br>(PDB 9PSI) |
| **Data collection and processing** |  |  |  |
| Magnification | 81,000 | 81,000 | 81,000 |
| Voltage (kV) | 300 | 300 | 300 |
| Electron exposure (e-/Å<sup>2</sup>) | 66 | 66 | 66 |
| Defocus range (μm) | 1.4-2.4 | 1.4-2.4 | 1.4-2.4 |
| Pixel size (Å) | 1.068 | 1.068 | 1.068 |
| Final Micrographs (no.) | 13,841 | 13,841 | 13,841 |
| Initial particle images (no.) | 6,208,655 | 6,208,655 | 6,208,655 |
| Final particle images (no.) | 20,794 | 32,749 | 27,408 |
| Symmetry imposed | C1 | C1 | C1 |
| Map resolution (Å) | 3.18 | 3.08 | 3.12 |
| FSC threshold | 0.143 | 0.143 | 0.143 |
| **Refinement** |  |  |  |
| Model resolution (Å) | 3.6 | 3.4 | 3.5 |
| FSC threshold | 0.5 | 0.5 | 0.5 |
| Model composition |  |  |  |
| Non-hydrogen atoms | 179,960 | 184,618 | 181,394 |
| Protein residues | 14,845 | 15,239 | 14,845 |
| Nucleotide | 2,760 | 2,836 | 2,828 |
| Ligands | 276 | 276 | 276 |
| *B* factors (Å<sup>2</sup>) |  |  |  |
| Protein | 192 | 181 | 182 |
| Nucleotide | 169 | 158 | 160 |
| Ligand | 146 | 129 | 136 |
| R.m.s. deviations |  |  |  |
| Bond lengths (Å) | 0.005 | 0.006 | 0.005 |
| Bond angles (°) | 0.8 | 0.9 | 0.8 |
| Validation |  |  |  |
| MolProbity score | 1.3 | 1.3 | 1.1 |
| Clashscore | 3.7 | 3.4 | 2.7 |
| Poor rotamers (%) | 0.00 | 0.01 | 0.02 |
| Ramachandran plot |  |  |  |
| Favored (%) | 97.49 | 97.33 | 97.68 |
| Allowed (%) | 2.49 | 2.64 | 2.31 |
| Disallowed (%) | 0.02 | 0.03 | 0.01 |

**Extended Data Table 5.** *TACO1* pathogenic mutations.

| Genetic variant | Protein | Clinical manifestations |
| --- | --- | --- |
| c.252_253delCT | p.Cys85PhefsTer15 | Childhood-onset progressive cerebellar and pyramidal syndrome with optic atrophy and learning difficulties<sup>19</sup>. |
| c.421C>T | p.Arg141Ter | Leigh syndrome due to mitochondrial complex IV deficiency<sup>18</sup>. |
| c.472insC | p.His158ProfsTer8 | Mitochondrial cytochrome c oxidase deficiency and late-onset Leigh syndrome<sup>22</sup>.<br>Childhood-onset progressive cerebellar and pyramidal syndrome with optic atrophy and learning difficulties<sup>19</sup>.<br>Slowly progressive cognitive dysfunction, dystonia or visual impairment<sup>17</sup>. |
| c.676G>T | p.Glu226Ter | Mitochondrial Complex IV deficiency. Adult-onset slowly progressive spastic paraparesis with cognitive impairment and a subcortical U-fiber leukoencephalopathy<sup>20</sup>. |
| c.777del | p.Glu259Aspfs*31 | Juvenile-onset Leigh syndrome<sup>21</sup>. |

**Extended Data Table 6.**
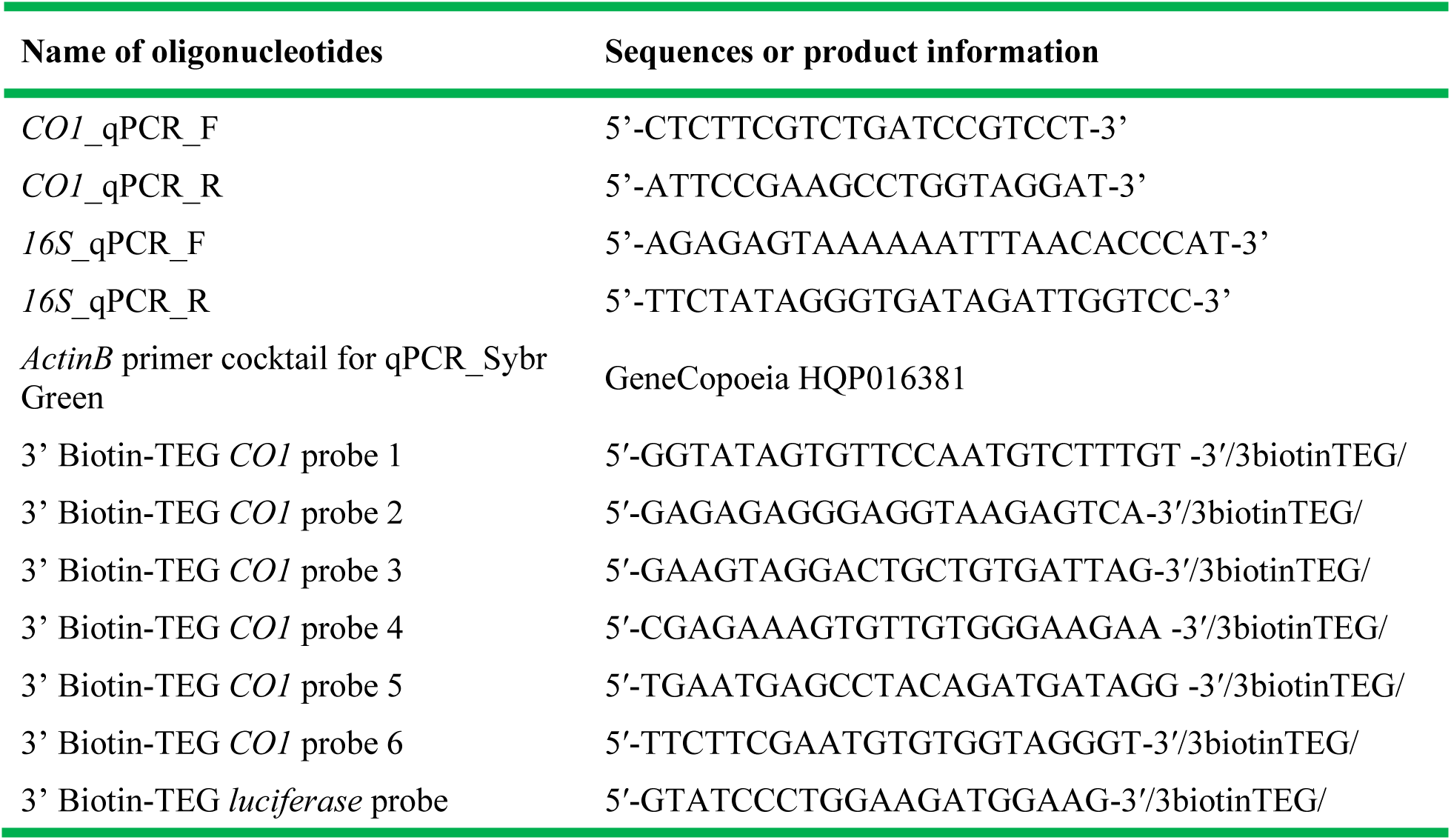
List of oligonucleotides used in the mRNA-specific binding protein assay.

